# Rewiring protein binding specificity in paralogous DRG/DFRP complexes

**DOI:** 10.1101/2023.05.31.543024

**Authors:** Christian A. E. Westrip, Stephen J Smerdon, Mathew L. Coleman

## Abstract

The Developmentally Regulated GTP-binding (DRG) proteins are an ancient subfamily of GTPases implicated in the regulation of translation and cell growth. In eukaryotes, there are two paralogs: DRG1 and DRG2, both of which have a conserved binding partner called DRG family regulatory protein 1 and 2 (DFRP1 and DFRP2), respectively. These binding partners are required for the function of DRGs, including their stabilisation at the protein level. Moreover, DFRPs interact with their respective DRG via a conserved region called the DFRP domain. Despite being highly similar, DRG1 and DRG2 have strict binding specificity for their respective DFRP. Using AlphaFold generated structure models of the human DRG/DFRP complexes, we have biochemically characterised their interactions and identified interface residues involved in determining specificity. This analysis revealed that as few as five mutations in DRG1 are able to switch its binding from DFRP1 to DFRP2. We show how two DRG1 residues in the core of the interface are most important for specifying the interaction with DFRP1 over DFRP2. We also demonstrate that whilst DFRP1 can stimulate the GTPase activity of DRG1, DFRP2 binding cannot. Overall, this work provides new insight into the structural determinants responsible for the binding specificities of the DRG:DFRP translation factor complexes, which are known to be essential for normal development in mice and humans.

## Introduction

Protein-protein interactions are an essential component of all biological processes. The manner in which proteins interact, often termed molecular recognition, is primarily determined by non-covalent interactions at the interface between proteins (Jones and Thornton, 1996; Keskin et al., 2008; Sheinerman et al., 2000). These interactions include hydrogen bonds, electrostatic interactions, van-der-Waals’ forces, and the hydrophobic effect (Jones and Thornton, 1996; Keskin et al., 2008; Sheinerman et al., 2000). The combination of these interactions at an interface, as well as the size of the interface, ultimately defines the binding affinity of an interaction (Kastritis and Bonvin, 2013; Keskin et al., 2008). Importantly, proteins are capable of specific binding, without which they would be unable to carry out defined functions. This binding specificity can be described as a protein interacting with a specific binding partner whilst *not* interacting with other proteins (Schreiber and Keating, 2011; Teilum et al., 2021). The formation of specific interactions is thought to be constrained by several factors. The inside of a cell is packed with millions of proteins, which often results in cognate interactors being saturated by non-cognate proteins. Many protein-protein interfaces also share similarities in shape with at least one other interface (Gao and Skolnick, 2010). This is largely due to the fact that most interfaces are more or less flat, suggesting that specificity must be achieved through other mechanisms besides shape complementarity. This problem is exemplified by the existence of many families of paralogous protein complexes, which due to the nature of paralogs share considerable sequence and structural similarities at their interfaces (Skerker et al., 2008). Despite these constraints, many paralogous proteins have achieved a remarkable level of specificity (Aakre et al., 2015; Capra et al., 2010; Fiebig et al., 2010; Skerker et al., 2008).

Specific binding is largely accomplished through non-covalent interactions at the interfaces of proteins (Aakre et al., 2015; Lite et al., 2020; Schreiber and Keating, 2011), though the temporal and spatial expression of a protein, together with its local concentration, may also play a role (Ivarsson and Jemth, 2019; Teilum et al., 2021). That is to say, two proteins need to be expressed at the same time and in the same place for them to interact. Conversely, expression at different times and in separate locations may help avoid an interaction. However, many thousands of proteins are expressed together in similar locations and at high concentrations, highlighting the importance of additional mechanisms to ensure biological specificity is achieved. The interface between two proteins typically contains residue-residue contacts that promote the interaction. These contacts are sometimes described as positive elements. Meanwhile, there may also exist residues in the interface that serve to block any non-cognate protein from binding and can be referred to as negative elements. These positive and negative elements are not necessarily mutually exclusive, as an interface residue can both promote one interaction whilst inhibiting another (Lite et al., 2020). As such, only a few interface residues may be needed to specify one interaction over another.

Whilst the general principles of protein-protein interactions, including binding specificity, have been well studied in some contexts, there still exist many individual interactions that remain poorly characterised. The Developmentally Regulated GTP-binding (DRG) proteins and their interactions with DRG Family Regulatory Proteins (DFRPs) represent one such example of protein binding specificity that has remained relatively unexplored. Eukaryotes typically have two paralogous DRG proteins, DRG1 and DRG2, both of which are highly conserved GTPases (Figure 1A) (Li and Trueb, 2000; Westrip et al., 2021). The potential importance of DRG proteins is exemplified by recent findings demonstrating that DRG1 knockout mice exhibit preweaning lethality, and that pathogenic DRG1 variants cause a novel neurodevelopmental disorder in humans (Westrip et al., 2023).

**Figure 1:**
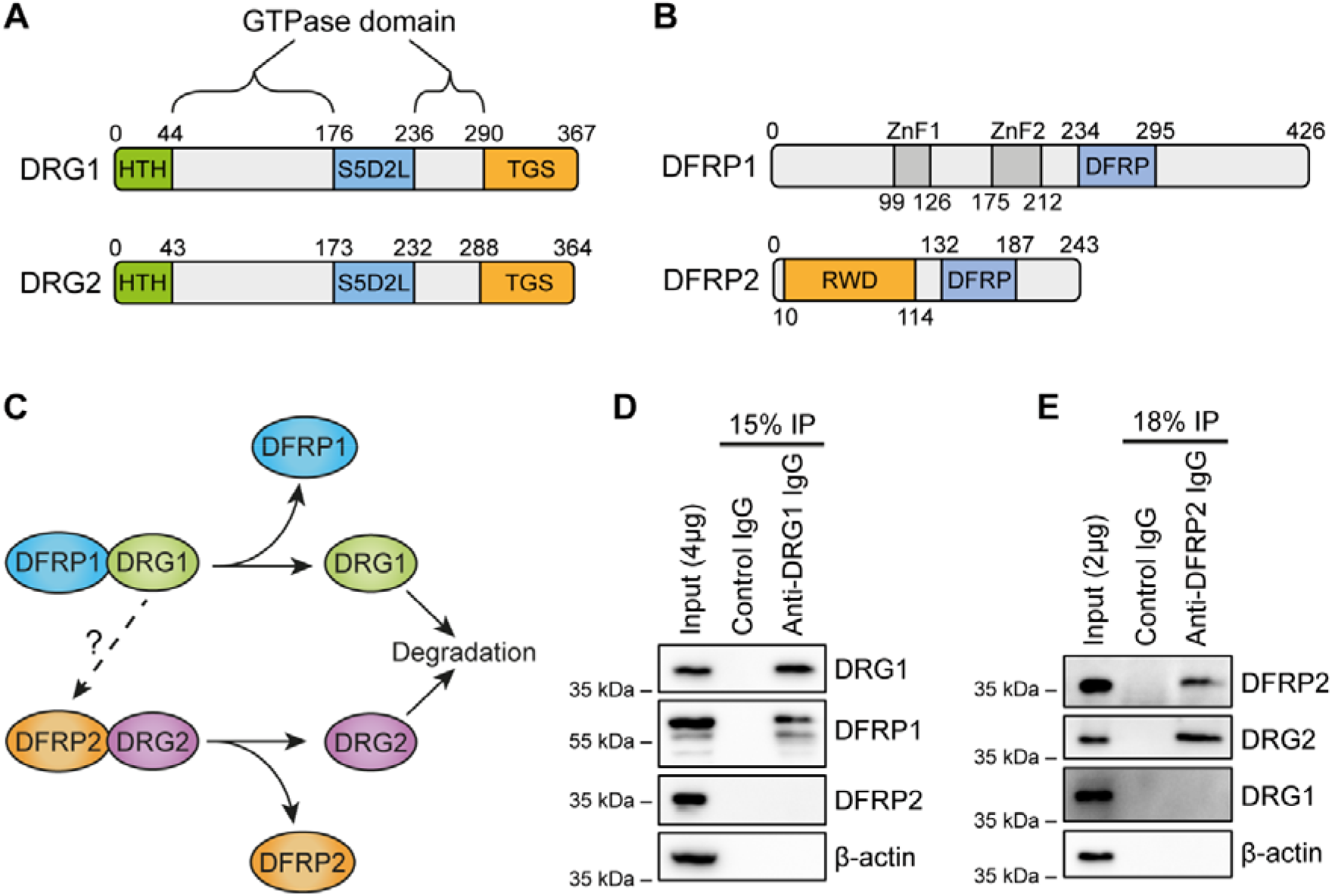
DRGs and DFRPs form specific complexes. **(A-B)** Domain architecture of the paralogous GTPases DRG1 and DRG2, and their respective binding partners DFRP1 and DFRP2. HTH: Helix-turn-Helix. S5D2L: Ribosomal protein S5 domain 2-like domain, TGS: ThrRS, GTPase, and SpoT domain. Znf: zinc finger. DFRP: DRG family regulatory protein domain. RWD: ring-finger containing proteins, <u>W</u>D-repeat-containing proteins, and yeast <u>D</u>EAD (DEXD)-like helicases domain. **(C)** Diagram illustrating the specific complexes formed: the DRG1/DFRP1 and the DRG2/DFRP2 complexes. Some weak binding may occur between DRG1 and DFRP2 when both are co-expressed. Removal of DFRP1/2 results in the degradation of the respective DRG protein. **(D)** Endogenous immunoprecipitation (IP) of the DRG1 protein from HEK293T cells followed by western blotting for the indicated proteins. **(E)** Endogenous IP of DFRP2 from HEK293T cells followed by western blotting for the indicated proteins.

Both DRG1 and DRG2 have a specific DFRP binding partner: DRG1 interacts with DFRP1, whilst DRG2 interacts with DFRP2 (Figures 1B and 1C) (Ishikawa et al., 2005). Although DRG1 and DRG2 are paralogs, their binding partners DFRP1 and DFRP2 are not. However, the DRG/DFRP interface is thought to be conserved between both complexes, as DFRP1 and DFRP2 share a conserved region called the DFRP domain that is required for the interaction with their respective DRG (Figure 1B) (Francis et al., 2012; Ishikawa et al., 2005). Moreover, both DRG/DFRP complexes are abundantly expressed in a variety of tissues and cell types and typically co-localise to the cytoplasm (Ishikawa et al., 2003; Li and Trueb, 2000; Westrip et al., 2021).

Although the function of these specific interactions is not fully understood, it is known that DFRP binding prevents the degradation of DRGs, likely via the ubiquitin-proteasome pathway (Figure 1C) (Ishikawa et al., 2005). As such, it is thought that the DRGs exist predominantly in complex with their DFRPs, as obligate heterodimers. The formation of these specific complexes is also likely to be important for their function in protein synthesis, as both the DRG1/DFRP1 and DRG2/DFRP2 complexes have been suggested to promote translation elongation (Kriachkov et al., 2023; Pochopien et al., 2021; Zeng et al., 2021). Work primarily performed in yeast has suggested that the DRG/DFRP complexes are recruited to stalled/slowly elongating ribosomes to promote efficient elongation. Notably, the DFRP proteins may play a role in directing DRGs to specific ribosome states (Daugeron et al., 2011; Pochopien et al., 2021; Zeng et al., 2021).

As mentioned above, DFRP1 and DFRP2 are not paralogs and aside from the DFRP domain, have entirely different domain organisations (Figure 1B). DFRP1 contains two zinc fingers whilst DFRP2 contains an RWD domain (Figure 1B). In humans, DFRP1 is over 180 amino acids bigger than DFRP2. In yeast, it has been suggested that DRG1 is recruited to translating ribosomes through its interaction with DFRP1 (Francis et al., 2012). This may rely on the zinc fingers of DFRP1 interacting with the ribosomal RNA of the small subunit (Zeng et al., 2021). Interestingly, DFRP2 may also be responsible for DRG2 recruitment to ribosomes, but through a different mechanism. The DRG2/DFRP2 complex was suggested to bind collided ribosomes in complex with Gcn1 (Pochopien et al., 2021). In particular, the RWD domain of DFRP2 may be required for this Gcn1 interaction (Ishikawa et al., 2013). As such, the DFRPs may be responsible for directing DRGs to related yet distinct pathways in the regulation of translation. Consistent with this theory, evidence in the literature suggests DRG1 and DRG2 are not functionally redundant, despite being highly similar (Westrip et al., 2021). If DFRP binding is responsible for the differing functions of DRG1 and DRG2, then a better understanding of their interaction specificity will aid in detailed characterisation of these important complexes.

Here, we explore the interaction mechanism between DRGs and DFRPs to identify the structural determinants of their binding specificity. Using AlphaFold generated structure models of the two human DRG/DFRP complexes we have identified interface residues that are responsible for DRG1 specifying binding to DFRP1 over DFRP2. Moreover, we demonstrate how it is possible to alter the specificity of DRG1 by switching its binding from DFRP1 to DFRP2. Our findings shed light on evolutionary constraints that drive protein binding specificity and raise questions regarding the role of DFRPs in regulating DRG GTPase activity.

## Results

### DRGs and DFRPs form specific complexes

We first confirmed the specific binding between DRGs and DFRPs using endogenous immunoprecipitation (IP) of the complexes from HEK293T cells. An endogenous IP of DRG1 resulted in co-purification of DFRP1 but not DFRP2 (Figure 1D). Purification of endogenous DFRP2 showed DRG2 but not DRG1 binding (Figure 1E). We also performed an endogenous DFRP1 IP, which showed enrichment for DRG1 (Figure S1A). Unfortunately, however, it was not possible to accurately assess DRG2 binding in the DFRP1 IP experiment (Figure S1A) due to cross-reactivity between anti-DRG2 antibodies and the DRG1 protein (Figure S1B). As such, we were not able to confirm whether endogenous DRG2 interacts exclusively with DFRP2 and not DFRP1. However, previous work has suggested DRG2 and DFRP1 do not interact *in vivo* (Ishikawa et al., 2005).

To further characterise the binding specificity, we co-transfected HEK293T cells with plasmids encoding HA-tagged DRG1 or DRG2, with or without V5-tagged DFRP1 or DFRP2, followed by an anti-HA IP and western blotting (Figure S1C). Although HA-DRG1 showed an abundant interaction with DFRP1-V5, it also showed some weak binding to DFRP2-V5, consistent with previously reported overexpression experiments (Ishikawa et al., 2005). HA-DRG2, however, interacted exclusively with V5-DFRP2 (Figure S1C). Together, the IP results with endogenous (Figures 1D and 1E) and exogenous (Figure S1C) complexes, confirms that DRGs and DFRPs form specific complexes, consistent with earlier reports in the literature (Ishikawa et al., 2009; Ishikawa et al., 2005; Ishikawa et al., 2013).

### Modelling the structures of the DRG/DFRP complexes using AlphaFold

To study the specific binding between DRGs and DFRPs in more detail, we chose to model the complexes using the AlphaFold2 algorithm (Jumper et al., 2021) implemented on the ColabFold server (Mirdita et al., 2022). Using this server, we built structures of the human DRG1/DFRP1 and DRG2/DFRP2 complexes (Figures 2A and 2B, respectively).

**Figure 2:**
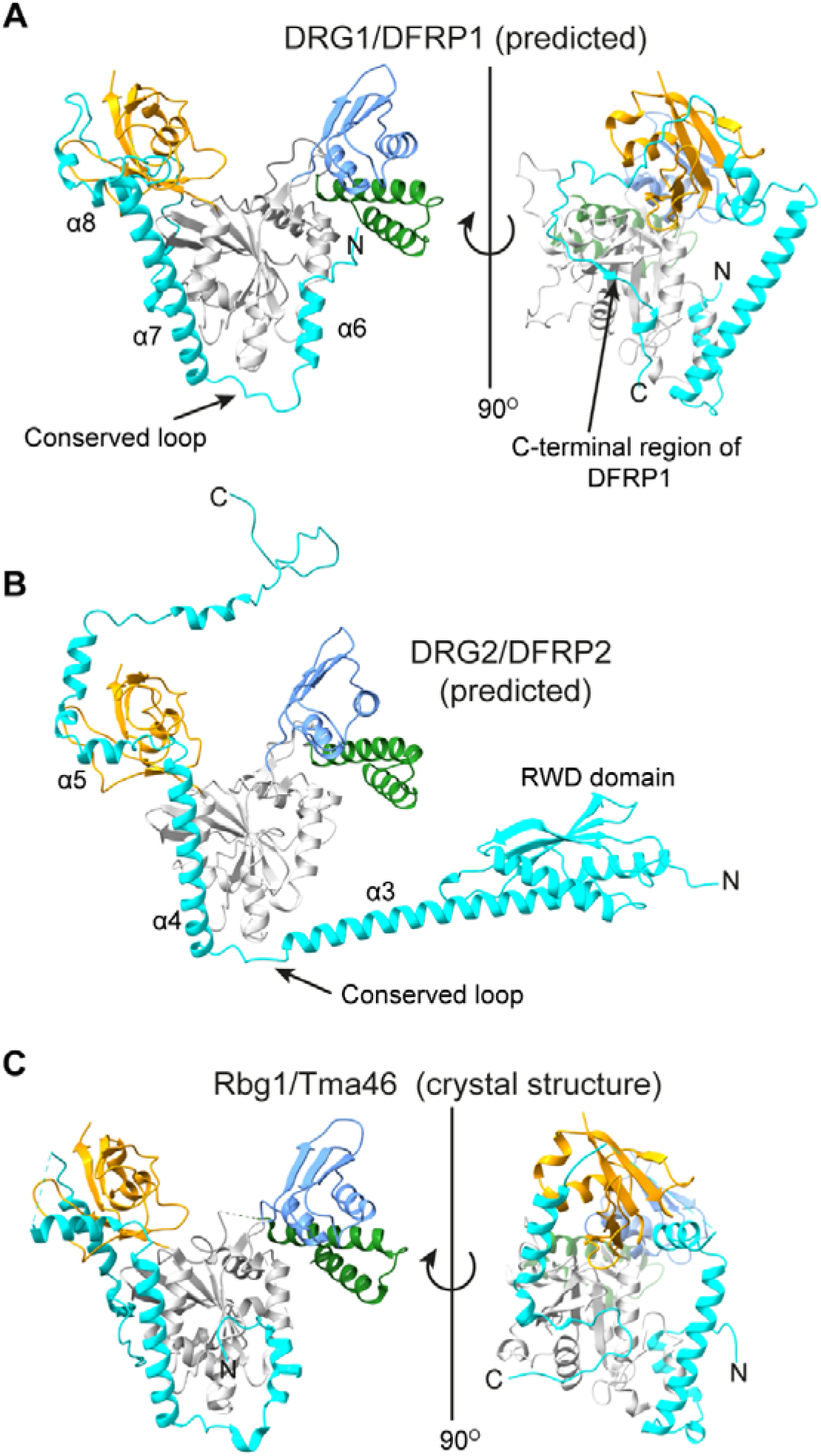
Predicted structural models of DRG/DFRP complexes. **(A)** Predicted structure of the human DRG1/DFRP1 complex. DFRP1 region K226-R340 is shown because the rest of the protein was poorly predicted. **(B)** Predicted structure of the human DRG2/DFRP2 complex. In both (A) and (B), DRG1/2 is coloured according to its domains. DFRP1 and DFRP2 are shown in cyan. **(C)** Crystal structure of the Rbg1/Tma46 (yeast DRG1/DFRP1) complex previously reported by Francis et al. (2012). Pdb: 4A9A. Rbg1 is coloured the same as in (A) and (B), whilst Tma46 is shown in cyan.

Both DRG1 and DRG2 were well predicted as they closely resembled the previously solved crystal structure of Rbg1 (yeast DRG1) (Figures S2A and S2B). The predicted structures show DRG1 and DRG2 have a typical GTPase domain, as well as the addition domains described earlier (Figure 1A).

Figure 2A reveals DFRP1 interacts with DRG1 via a series of helices and loops that wrap around DRG1, contacting both the GTPase and TGS domains (Figure 2A). This is very similar to the previously reported crystal structure of the orthologous Rbg1/Tma46 (yeast DRG1/DFRP1) complex (Figure 2C) (Francis et al., 2012). Structural alignments between the predicted DRG1/DFRP1 model and the Rbg1/Tma46 crystal structure further highlighted the similarities (Figures S2A and S2C). It should be noted that DFRP1 contains large regions of disorder in both the N- and C-termini that were poorly predicted by AlphaFold2 and so are not shown in Figure 2A.

DFRP2 contacted DRG2 in a largely similar manner to DFRP1 and DRG1 (Figures 2A and 2B). Structural comparison with the Rbg1/Tma46 crystal structure also revealed similarities with the DRG2/DFRP2 complex (Figures S2B and S2D). Both DFRP1 and DFRP2 contact their respective DRG through a conserved loop, followed by a long helix (α7 and α4 in DFRP1 and DFRP2, respectively) and then a short helix (α8 and α5 in DFRP1 and DFRP2, respectively) (Figures 2A and 2B).

Despite these similarities, there were also differences between the two predicted DRG/DFRP complexes. DFRP1 forms additional contacts with DRG1, through a C-terminal region that wraps around the TGS domain and forms further interactions with the GTPase domain (Figure 2A). This closely resembles the manner in which Tma46 interacts with Rbg1 in the crystal structure (Figures 2C and S2A). On the other hand, DFRP2 does not make these contacts with DRG2, with the DFRP2 C-terminal region instead predicted as a disordered tail (Figure 2B).

DFRP1 also has a short helix (α6) that inserts into a pocket on DRG1, similar to the Rbg1/Tma46 crystal structure (Figures 2A and 2C). This short helix was absent in DFRP2, and instead there is a long alpha helix (α3) that connects to the RWD domain (Figure 2B).

Whilst DFRP1 had large regions of predicted disorder, including the zinc fingers, DFRP2 was more ordered, especially in its N-terminal RWD domain (Figure 2B). The RWD domain of DFRP2 was well predicted as shown by its high similarity to the already solved NMR structure of this domain (Figure S2E) (Yokoyama et al., 2008). Overall, the comparative structural analyses suggested the predicted structural complexes were appropriate for further investigation.

### Identifying interface residues in DRG/DFRP complexes

Using the predicted structure models (Figure 2), we identified the residues that are located in the interfaces between DRG1 and DFRP1, and between DRG2 and DFRP2. Analysis of the predicted DRG1/DFRP1 complex revealed a large number of contacts: 60 residues in DRG1 contacted 45 residues in DFRP1 (Table S1). Meanwhile, analysis of the predicted DRG2/DFRP2 complex identified 35 residues in DRG2 contacting 29 residues in DFRP2 (Table S2). The buried surface area for the DRG1/DFRP1 interface was 2988 Å, whilst for the DRG2/DFRP2 complex it was 1697 Å. The larger interface of the DRG1/DFRP1 complex is due to the fact that DFRP1 wraps around DRG1 more extensively than DFRP2, contacting the TGS domain from opposite sides, consistent with the Rbg1/Tma46 crystal structure as discussed above (Figure 2). Identified interface residues were mapped onto protein sequence alignments of DRGs and DFRPs (Figures 3A and 3B) (see also Figures S3 and S4 for full alignments). The DRG1/2 alignment shows that there are a significant number of shared positions in DRG1 and DRG2 that both contact their respective DFRP (blue triangles) (Figures 3A and S3). There were also numerous residues that only featured in the DRG1/DFRP1 interface (green triangles). In contrast, there were few residues that are specific to the DRG2/DFRP2 interface (orange triangles) (Figures 3A and S4).

**Figure 3:**
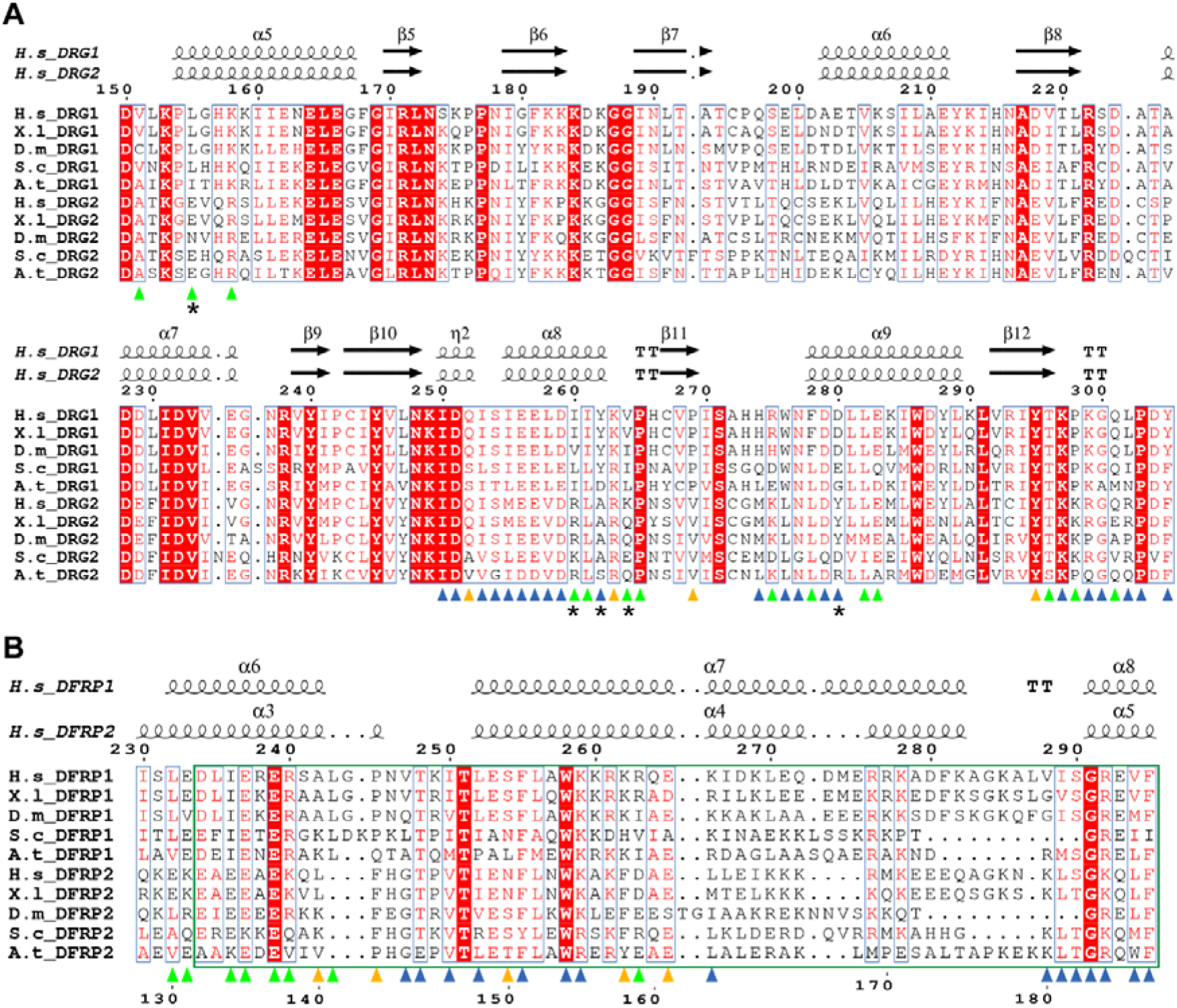
DRG and DFRP protein sequence alignments. **(A)** Alignment of DRG protein sequences. Asterisks mark positions in DRG1/2 that were investigated for their roles in binding specificity. **(B)** Alignment of DFRP protein sequence alignments. In both (A) and (B) the green arrows represent interface residues in the DRG1/DFRP1 complex, orange arrows for the DRG2/DFRP2 complex and blue arrows for interface residues present in both complexes. Secondary structure from the predicted human structures is shown above the alignments. Green box denotes the DFRP domain. See Supplemental Information for full alignments and a table of sequences used.

The partial DFRP1/2 protein sequence alignment in Figure 3B shows the conserved region required for DRG binding, previously named the DFRP domain (Ishikawa et al., 2005). Based on sequence analysis this region was previously defined as amino acids 234-295 in DFRP1 and 132-187 in DFRP2 (Ishikawa et al., 2005) (Figure 3B). Mapping the interface residues onto the alignment revealed there are several positions in DFRP1 and DFRP2 that are conserved between the two interfaces (Figure 3B, blue triangles). However, there are also a number of residues that are only present in one of the two interfaces (Figure 3B).

Further inspection of the DFRP domain shows the most conserved region is at the beginning of the long alpha helix (α7 DFRP1/α4 DFRP2) (Figure 3B). This region also includes multiple residues that contact DRG1 and DRG2. Following this is a short alpha helix (α8 DFRP1/α5 DFRP2) that also contains some highly conserved residues that contact both DRG1 and DRG2 (Figure 3B). There are also multiple interface residues in the region 300-340 in DFRP1 and 190-210 in DFRP2 (Figure S4). These regions are not conserved between DFRP1 and DFRP2. Interestingly, previous studies have shown that this C-terminal region of DFRPs is not required for DRG binding in either yeast or mammalian complexes (Francis et al., 2012; Ishikawa et al., 2005). Therefore, although the predicted structures suggested these regions of DFRP1/2 are part of the interfaces, they may not be essential for DRG binding.

### Conserved residue contacts between DRGs and DFRPs

The results in Figures 2 and 3 suggested the interfaces between DRGs and DFRPs are largely similar for both complexes, including numerous conserved residue contacts. We decided to investigate some of these residues for their requirement for complex formation. Previously, it had been shown that the Rbg1/Tma46 interface contains a number of aromatic residues forming potential pi-pi interactions, and that these contacts may be important for complex formation (Francis et al., 2012). Two of these aromatic residues occur in the DFRP domain and are conserved between DFRP1 and DFRP2: F255^DFRP1^/F151^DFRP2^ and W258^DFRP1^/W154^DFRP2^ (Figure 3B). Closer analysis of the tryptophan W258^DFRP1^/W154^DFRP2^ suggests it is invariant (Figure 3B) and makes numerous contacts with DRG1/2, respectively (Figures S5A and S5B). We tested the importance of this tryptophan by mutating it to an alanine in our V5-tagged DFRP1/2 vectors, followed by transient expression in HEK293T cells and an anti-V5 IP. Unsurprisingly, the W258A^DFRP1^ and W154A^DFRP2^ mutations significantly reduced binding to DRG1 and DRG2, respectively (Figures S5C and S5D).

We also analysed some of the highly conserved residues in DRG1 and DRG2 for their role in DFRP1/2 binding. Both DRGs have a highly conserved aspartate (D259^DRG1^ and D257^DRG2^) that contacts the DFRP1/2 tryptophan described above, potentially forming a hydrogen bond (Figure S6A). We also identified another aspartate (D279^DRG1^ and D277^DRG2^) that may form an electrostatic contact with a lysine residue in DFRP1/2 (K259^DFRP1^ and K155^DFRP2^, respectively) (Figure S6B). In addition, we also identified a tryptophan residue in DRG1 (W276^DRG1^) that may be involved in the pi-pi interactions with DFRP1, as mentioned above (Francis et al., 2012) (Figure S6C). We mutated these residues to alanines in HA-DRG1 and HA-DRG2 vectors, expressed them in HEK293T cells followed by an anti-HA IP and western blotting. Individual mutation of either of the two aspartates in DRG1 (D259A^DRG1^ and D279A^DRG1^) did not cause a significant reduction in DFRP1 binding (Figure S6D). However, a double mutation (D259A^DRG1^ + D279A^DRG1^) did cause a reduction in DFRP1 binding compared to wildtype DRG1. More strikingly, mutation of the DRG1 tryptophan (W276A^DRG1^) resulted in a significant loss of DFRP1 binding (Figure S6D).

In DRG2, the D257A^DRG2^ mutation caused a slight reduction in DFRP2 binding compared to wildtype DRG2, whilst D277A^DRG2^ did not (Figure S6E). Similar to the double aspartate mutation in DRG1, the combination of the mutations in DRG2 (D257A^DRG2^ + D277A^DRG2^) caused a larger loss of DFRP2 binding compared to the single D257A^DRG2^ mutation (Figure S6E). Together, the results in Figures S5 and S6 demonstrate how the predicted structure models have helped identify interface residue important for complex formation, thus supporting the accuracy of these structures.

### Identification of possible specificity determining residues

After establishing the predicted structure models were likely accurate, we chose to investigate which residues in DRG1 and DRG2 may be important for specifying binding to their respective DFRP. We hypothesised that these DRG1/2 residues would be in the interface but conserved *differently* between DRG1 and DRG2, as have been demonstrated for other paralogous complexes that show binding specificity (Aakre et al., 2015; Lite et al., 2020). For instance, an interface residue might be expected to control binding specificity if it is positively charged in DRG1 but negatively charged in DRG2. As such, we searched the DRG protein sequence alignment (Figures 3A and S3) for positions in the interface where the amino acid is significantly different in charge, size, or hydrophobicity between DRG1 and DRG2. As mentioned above, DFRP1 has a region C-terminal to the DFRP domain that wraps around DRG1 (Figures 2A and S2C). This region, however, is not well conserved between DFRP1 orthologs (Figure S4). Moreover, as this C-terminal region of DFRP1 may not be required for the interaction with DRG1 (Francis et al., 2012; Ishikawa et al., 2005), we decided to focus on DRG1/2 residues that contact the conserved DFRP domain of DFRP1/2 in our search for specificity determining residues.

Our analysis identified five sites that show, at these positions, considerably different amino acids between DRG1 and DRG2 (Figures 4A and 4B) (these positions are also labelled in the alignment in Figure 3A with asterisks). Three of the positions are conserved as hydrophobic amino acids in DRG1 (L155^DRG1^, I260^DRG1^, V264^DRG1^) whilst they are charged residues in DRG2 (E153^DRG2^, R258^DRG2^, K262^DRG2^). Another position of interest shows a change from a large (Y262^DRG1^) to a small residue (A260^DRG2^). The fifth position we identified is a charged residue in DRG1 (D280^DRG1^) whilst it is a tyrosine in DRG2 (Y278^DRG2^).

**Figure 4:**
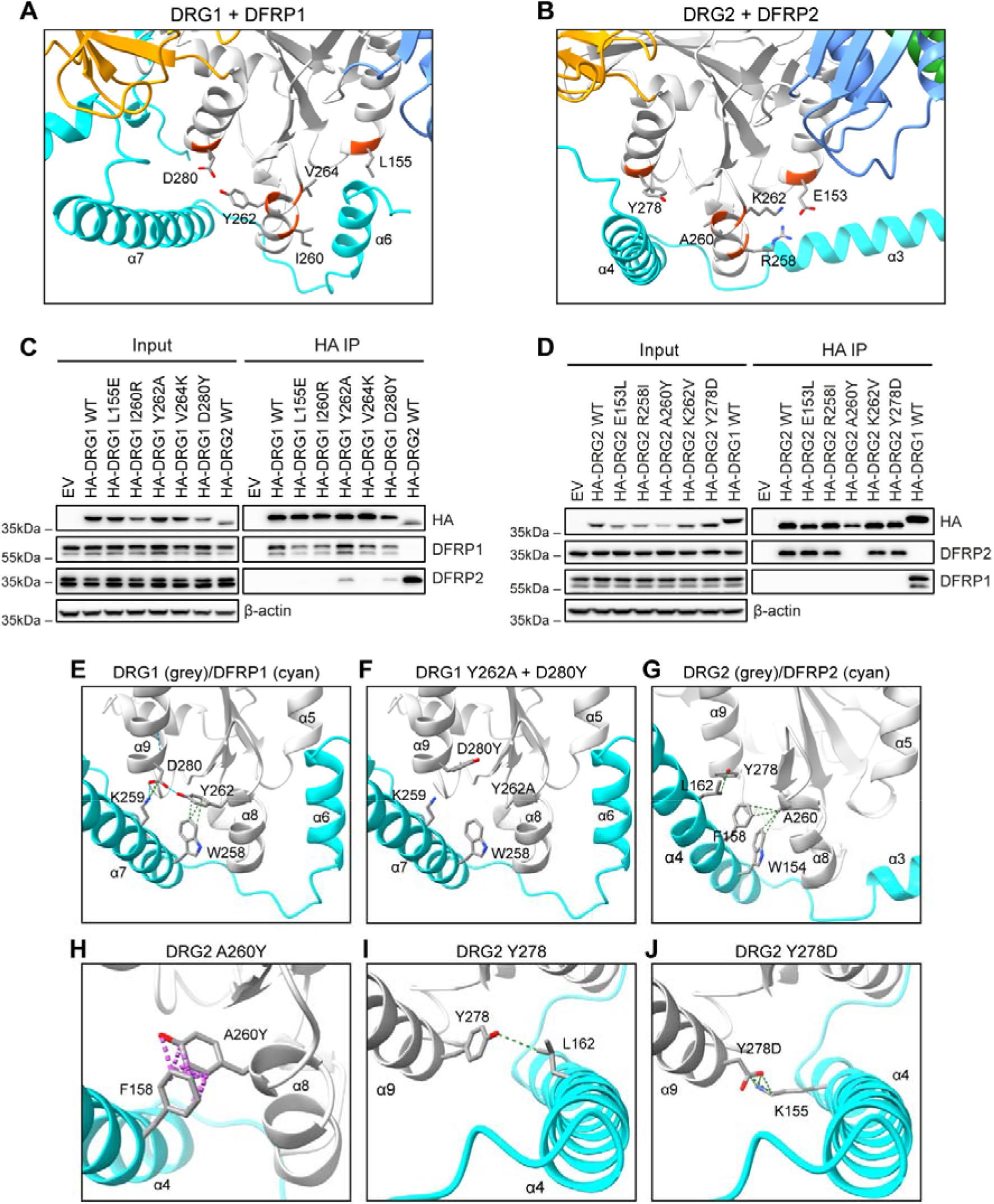
DRG1/2 residues involved in DFRP binding specificity. **(A)** Location of the five DRG1 amino acids we identified as possible DFRP specificity determining residues (main chain for these sites is coloured in red). **(B)** Same as in (A) but showing the corresponding positions in DRG2. **(C)** The DRG1 positions from (A) mutated to the corresponding residue in DRG2. Mutants were encoded in a HA-DRG1 vector and expressed in HEK293T cells followed by an anti-HA immunoprecipitation (IP). EV: empty vector plasmid. **(D)** Same as in (C) but with the reverse mutations encoded in a HA-DRG2 vector. **(E)** Contacts between Y262^DRG1^, D280^DRG1^ and the DFRP1 residues W258^DFRP1^ and K259^DFRP1^. **(F)** Structural modelling of the Y262A^DRG1^ and D280Y^DRG1^ mutations in DRG1. **(G)** Contacts between A260^DRG2^, Y278^DRG2^ and the DFRP2 residues W154^DFRP2^, F158^DFRP2^ and L262^DFRP2^. **(H)** Likely clashes between A260Y^DRG2^ mutation and F158^DFRP2^. **(I)** DRG2 residue Y278^DRG2^ contacts L162^DFRP2^. **(J)** The Y278D^DRG2^ mutation may contact K155^DFRP2^.

We hypothesised that mutating these positions in one paralog to the corresponding residue in the other paralog may switch DFRP binding. To test this, we first individually mutated each of the five positions in HA-DRG1 vectors to their respective positions in DRG2 (L155E^DRG1^, I260R^DRG1^, Y262A^DRG1^, V264K^DRG1^, D280Y^DRG1^). The HA-DRG1 mutants were expressed in HEK293T cells followed by an anti-HA IP (Figure 4C). Each of the three hydrophobic to charged residue mutations in DRG1 (L155E^DRG1^, I260R^DRG1^, V264K^DRG1^) resulted in a very small decrease in DFRP1 binding compared to wildtype DRG1, but did not cause any detectable DFRP2 binding (Figure 4C). Interestingly, both the Y262A^DRG1^ and D280Y^DRG1^ mutations resulted in noticeable DFRP2 binding, though not to the same levels as the HA-DRG2 wildtype control (Figure 4C). The D280Y^DRG1^ mutation may have also decreased DFRP1 binding relative to wildtype DRG1, whilst the Y262A^DRG1^ mutation did not.

In the same manner, we next tested the five reverse mutations (E153L^DRG2^, R258I^DRG2^, A260Y^DRG2^, K262V^DRG2^, Y278D^DRG2^) encoded in HA-DRG2 vectors for their effect on DFRP1/2 binding (Figure 4D). Strikingly, the A260Y^DRG2^ mutation resulted in a complete loss of DFRP2 binding compared to wildtype HA-DRG2, whilst the other four mutations had no effect on this interaction (Figure 4D). Surprisingly, none of the mutations, including A260Y^DRG2^, resulted in any detectable DFRP1 interaction, suggesting these mutations are not sufficient to switch DFRP binding (Figure 4D).

### Structural analysis of Y262^DRG1^, D280^DRG1^, A260^DRG2^, and Y278^DRG2^

Of the residues identified in DRG1 the Y262^DRG1^ and D280^DRG1^ positions appeared to be most important for specifying DFRP1 over DFRP2 (Figure 4C). Closer analysis of the Y262^DRG1^ residue in the predicted DRG1/DFRP1 complex, suggests it interacts with W258^DFRP1^ (Figure 4E) as part of the pi-pi interactions previously reported (Francis et al., 2012). The D280^DRG1^ residue may form a salt bridge with K259^DFRP1^. Interestingly, Y262^DRG1^ and D280^DRG1^ may also form a hydrogen bond with each other (Figure 4E). Modelling the mutations Y262A^DRG1^ and D280Y^DRG1^ suggested they would not cause any clashes with DFRP1, though the contacts described above would be lost (Figure 4F).

The corresponding positions in DRG2, A260^DRG2^ and Y278^DRG2^, were also further analysed. The A260^DRG2^ residue interacts with two aromatic amino acids in DFRP2, namely W154^DFRP2^ and F158^DFRP2^ (Figure 4G). Modelling the A260Y^DRG2^ mutation explains why this change disrupted DFRP2 binding (Figure 4D): The presence of the much larger tyrosine at residue A260^DRG2^ would likely cause a clash with the DFRP2 phenylalanine F158^DFRP2^ (Figure 4H). This may also explain why the reverse mutation in DRG1 (Y262A^DRG1^) allowed some DFRP2 binding to occur (Figure 4C), as this tyrosine in DRG1 (Y262^DRG1^) would presumably clash with the phenylalanine in DFRP2, thus preventing the DRG1-DFRP2 interaction. Consistent with this, mutation of the tyrosine Y262^DRG1^ to an alanine removes any potential clash with DFRP2, which could help maintain the interaction, as observed in Figure 4C.

In the DRG2/DFRP2 interface, Y278^DRG2^ contacts L162^DFRP2^ (Figure 4I). Modelling the mutation Y278D^DRG2^ suggests that whilst the L162^DFRP2^ contact may be lost, an interaction with a lysine, K155^DFRP2^, could be created (Figure 4J), potentially explaining the lack of effect on the DRG2:DFRP2 interaction (Figure 4D).

### DFRP1 interacts with a hydrophobic pocket in DRG1

The three other positions of interest in DRG1 were hydrophobic residues that are conserved as charged residues in DRG2 (Figures 3A, 4A and 4B). Interestingly, these three positions in DRG1 (L155^DRG1^, I260^DRG1^ and V264^DRG1^) form part of a hydrophobic pocket that DFRP1 inserts a short helix into (α6) (Figures 5A and 5B). The same region of DRG2 does not have this hydrophobic pocket owing to the presence of larger and charged residues compared to DRG1 (Figure 5C). As such, mutating the hydrophobic residues in DRG1 to charged amino acids (L155E^DRG1^, I260R^DRG1^ and V264K^DRG1^) would likely cause clashes with the short helix (α6) in DFRP1. This may explain why individual mutations of these DRG1 residues appeared to modestly reduce DFRP1 binding (Figure 4C). We hypothesise that the presence of charged residues at this position in DRG2 (Figure 5C) may function to prevent DFRP1 binding.

**Figure 5:**
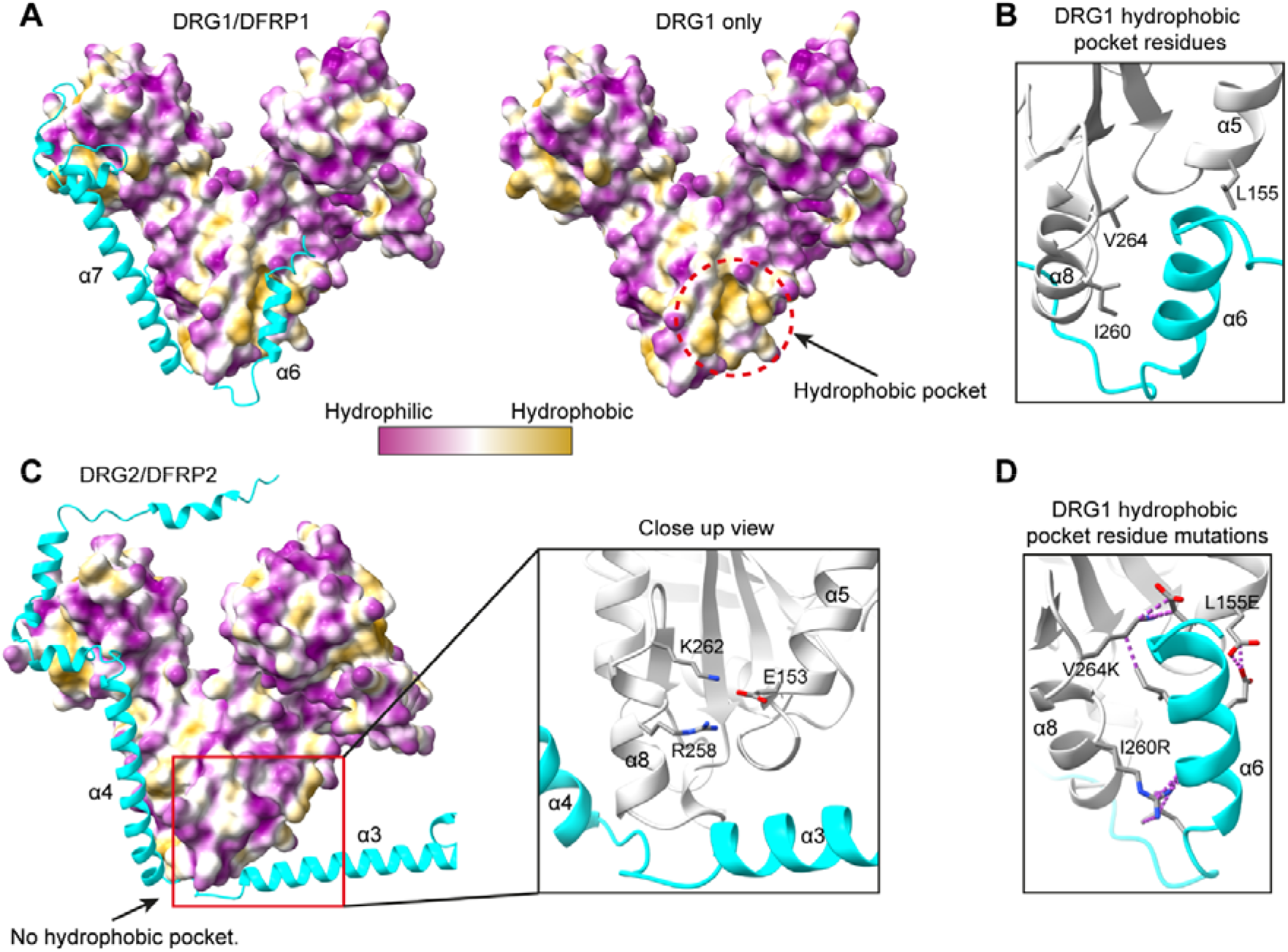
DFRP1 interacts with a hydrophobic pocket in DRG1. **(A)** Left: DRG1/DFRP1 predicted structure complex showing the surface representation of DRG1 coloured according to hydrophobicity. Right: without DFRP1. **(B)** Close-up view of the hydrophobic pocket, including the residues identified earlier as potential specificity determining residues. **(C)** The predicted DRG2/DFRP2 complex showing the surface representation of DRG2 coloured according to hydrophobicity, same as in (A). **(D)** Modelling the hydrophobic to charged residue mutations in DRG1, showing the potential clashes with DFRP1 as dashed purple lines.

### Combined mutations in DRG1 can switch specificity

We next tested whether combining the DRG1 mutations described above (Figure 4C) would result in a larger switch in DFRP1/2 binding compared to the individual mutations. As such, we created a HA-DRG1 vector encoding both the Y262A^DRG1^ and D280Y^DRG1^ mutations, which individually can modestly support DFRP2 binding (Figure 4C). We also created vectors that encode Y262A^DRG1^ and D280Y^DRG1^ mutations in combination with mutations in the hydrophobic pocket. For simplicity, we describe these mutant combinations by their single amino acid letter in DRG1, i.e., combined Y262A^DRG1^ + D280Y^DRG1^ mutations are referred to as YD^DRG1^.

Combination of the Y262A^DRG1^ and D280Y^DRG1^ (YD^DRG1^) mutations resulted in a decrease in DFRP1 binding compared to wildtype DRG1 and a significant amount of DFRP2 binding (Figure 6A). Addition of the hydrophobic pocket mutation L155E^DRG1^ to the YD^DRG1^ mutant (LYD^DRG1^) resulted in a larger loss of DFRP1 binding compared to wildtype DRG1 (Figure 6A). Furthermore, the LYD^DRG1^ mutation showed a further increase in DFRP2 binding relative to YD^DRG1^. Addition of the other two hydrophobic pocket mutations, I260R^DRG1^ and V264K^DRG1^, to LYD^DRG1^ (creating the LIYD^DRG1^ and LIYVD^DRG1^ mutant combinations) only modestly reduced DFRP1 binding further (Figure 6A).

**Figure 6:**
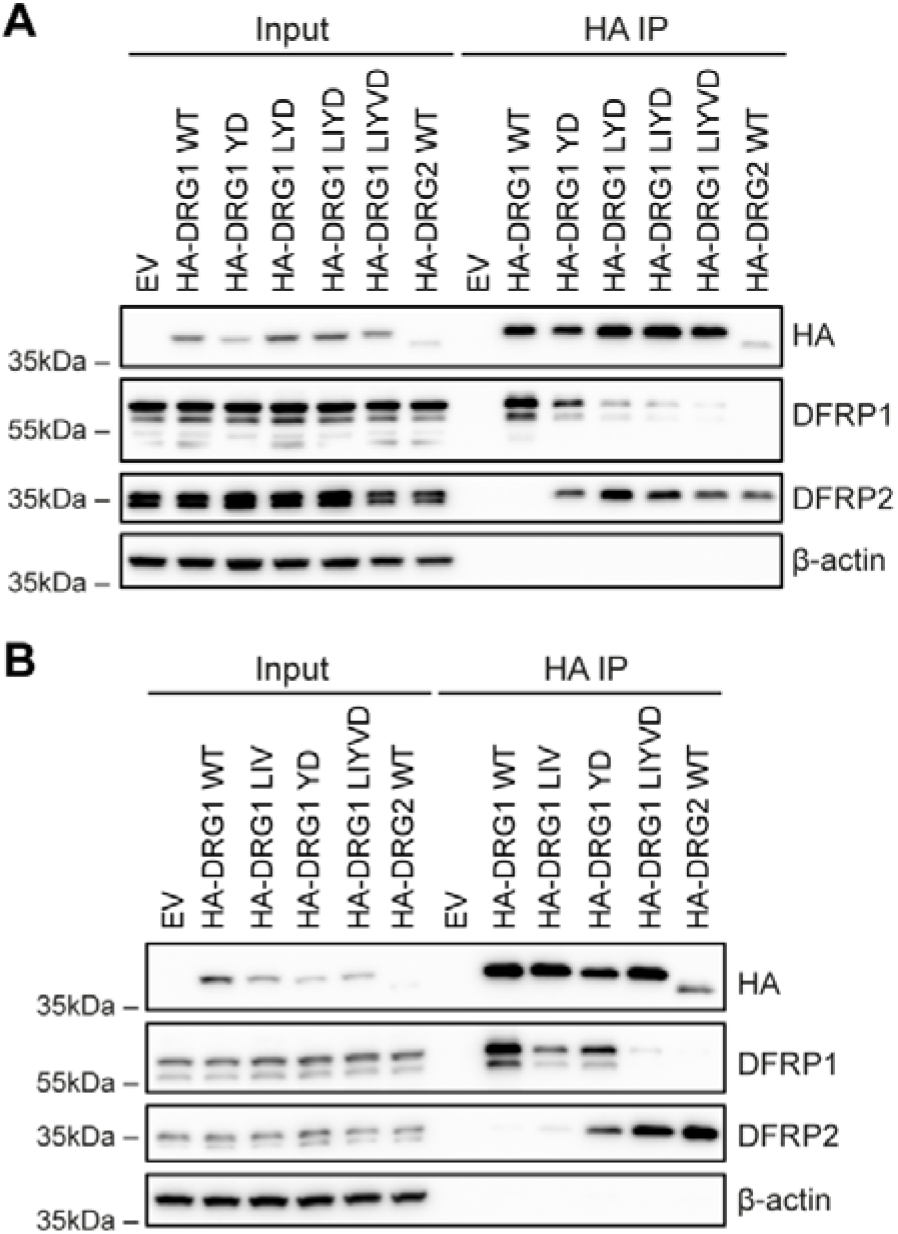
Combined DRG1 mutations switch binding from DFRP1 to DFRP2. **(A)** HEK293T cells were transfected with vectors encoding HA-DRG1 wildtype (WT) or the indicated combination of mutations: L: L155E^DRG1^, I: I260R^DRG1^ Y: Y262A^DRG1^, V: V264K^DRG1^, D: D280Y^DRG1^. HA-DRG2 WT was also included as a control for DFRP2 binding. The HEK293T cell lysates were used in an anti-HA immunoprecipitation (IP) followed by western blotting for the indicated proteins. **(B)** Same as in (A) but with the L155E^DRG1^ + I260R^DRG1^ + V264K^DRG1^: LIV^DRG1^ vector included. EV: empty vector plasmid.

The results in Figure 6A raised the possibility that the loss of DFRP1 binding, through mutations in the hydrophobic pocket, could allow a DFRP2 interaction, even though these residues are not predicted to contact DFRP2 (see Figure 5). To further dissect the role of these residues, we created a HA-DRG1 vector encoding all three hydrophobic pocket mutations in isolation (L155E^DRG1^ + I260R^DRG1^ + V264K^DRG1^: LIV^DRG1^) and compared its DFRP1/2 binding to the YD^DRG1^ and LIYVD^DRG1^ vectors in an anti-HA IP experiment (Figure 6B).

Mutating all three hydrophobic residues (LIV^DRG1^) resulted in a significant loss of DFRP1 binding compared to wildtype DRG1, whilst only producing a very small amount of DFRP2 binding (Figure 6B). Consistent with the earlier result in Figure 6A, the YD^DRG1^ mutation resulted in considerable DFRP2 binding whilst also reducing DFRP1 binding (Figure 6B). Importantly, whilst the LIV^DRG1^ mutant resulted in some DFRP2 binding, it was considerably less than the YD^DRG1^ vector, despite the LIV^DRG1^ mutant causing a greater loss in DFRP1 binding. This suggest that loss of DFRP1 binding alone is not sufficient to switch the interaction to DFRP2. Instead, the YD^DRG1^ mutations appear to be needed to specify DFRP2 binding. Thus, when LIV^DRG1^ and YD^DRG1^ are combined to create LIYVD^DRG1^, we observed a more complete switch in binding from DFRP1 to DFRP2 (Figure 6B).

### DRG1 LIYVD shows promiscuous binding for co-transfected DFRP1 and DFRP2

In Figure 6, we demonstrated how mutation at five sites in DRG1 can switch binding from DFRP1 to DFRP2. However, what is not clear from these experiments is whether the LIYVD^DRG1^ mutant binds DFRP2 to the same extent as HA-DRG2 does. This is due to the fact that HA-DRG2 was considerably underexpressed relative to HA-DRG1, resulting in HA-DRG2 not being enriched to comparable levels in the HA IP (Figure 6). We therefore chose to test the HA-DRG1 LIYVD mutant in an HA IP with co-transfection of V5-tagged DFRP1 or DFRP2 (Figure 7). Consistent with the result in Figure S1C, HA-DRG1 interacted abundantly with DFRP1-V5 and only showed weak binding to DFRP2-V5 (Figure 7). Interestingly, LIYVD^DRG1^ was able to interact with DFRP1-V5 when overexpressed together, although the binding was reduced compared to wildtype HA-DRG1 (Figure 7, compare lanes 17 and 20). However, LIYVD^DRG1^ and DFRP2-V5 also interacted abundantly when overexpressed together. Furthermore, this interaction was comparable to the HA-DRG2/DFRP2-V5 interaction (Figure 7, compare lanes 21 and 24). These results suggest that DRG1 LIYVD has promiscuous binding in that it can interact with either DFRP1 or DFRP2.

**Figure 7:**
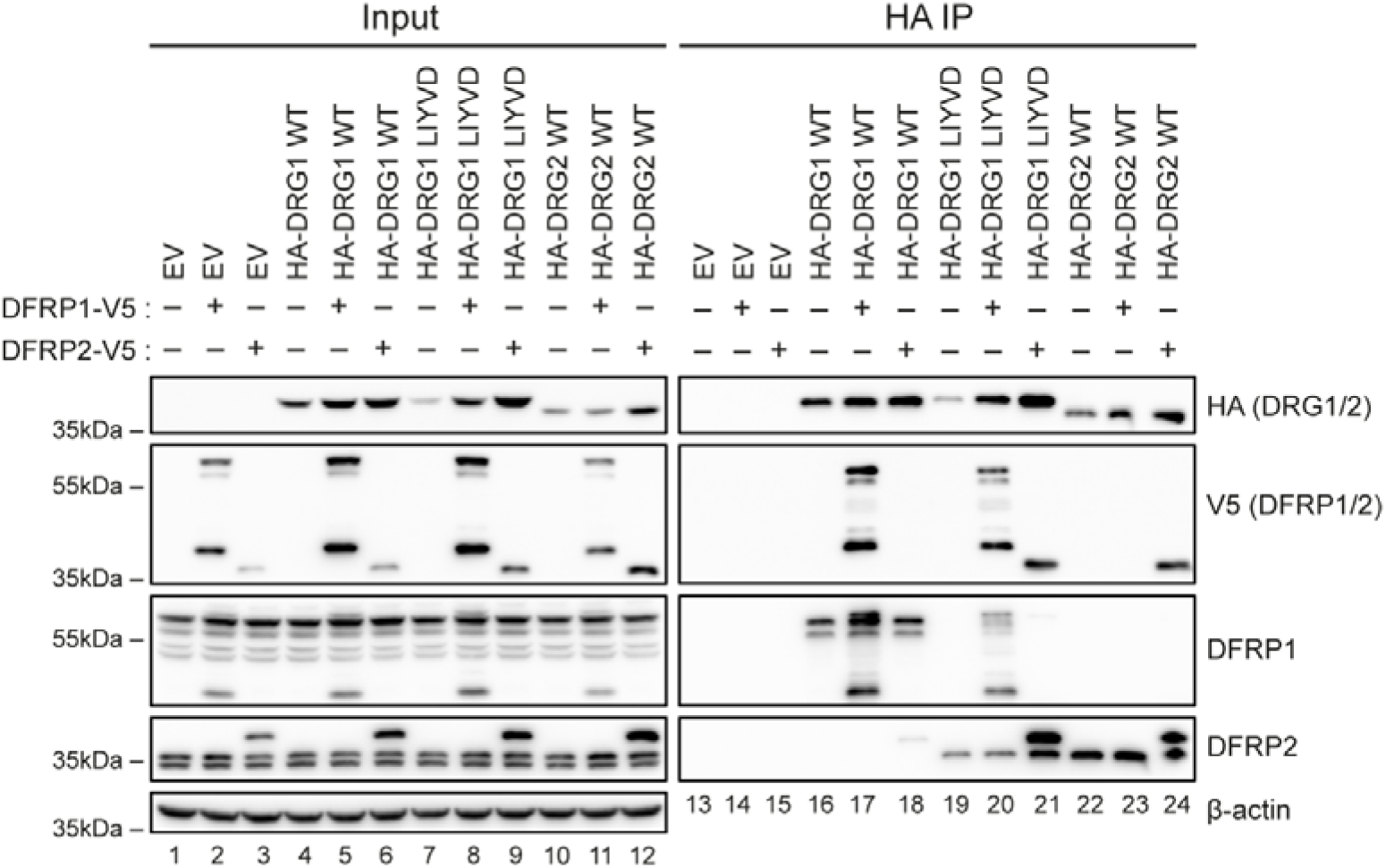
DRG1 LIYVD has promiscuous binding depending on DFRP availability. HEK293T cells were co-transfected with HA-DRG1/2 vectors with or without DFRP1-V5 or DFRP2-V5. Cell lysates were used in an anti-HA immunoprecipitation (IP) followed by western blotting. LIYVD: L155E^DRG1^ + I260R^DRG1^ + Y262A^DRG1^ + V264K^DRG1^ + D280Y^DRG1^. EV: empty vector plasmid. WT: wildtype.

### Combined mutations in DRG2 do not switch its specificity

Considering it was possible to alter the specificity of DRG1 with the five mutations described above (see Figure 6), we expected the combined reverse mutations in DRG2 would also changes its DFRP binding specificity (i.e., switch its binding from DFRP2 to DFRP1). Therefore, we created HA-DRG2 vectors encoding combinations of the mutations described in Figure 4D, as was performed for DRG1 (see Figure 6A). These HA-DRG2 mutants were expressed in HEK293T cells followed by an anti-HA IP and western blotting (Figure S7A). Surprisingly, combining A260Y^DRG2^ and Y278D^DRG2^ mutations (AY^DRG2^) resulted in no detectable DFRP1 binding, despite loss of the DFRP2 interaction (Figure S7A). Moreover, addition of the other DRG2 mutations (E153L^DRG2^, R258I^DRG2^, K262V^DRG2^, creating EAY^DRG2^, ERAY^DRG2^ and ERAKY^DRG2^) failed to switch binding to DFRP1, also despite the loss of DFRP2 binding (loss of DFRP2 binding is likely due to the A260Y^DRG2^ mutation alone, as described earlier; Figures 4D and 4H).

One possible reason for the inability of ERAKY^DRG2^ to interact with DFRP1 may stem from the differences in the C-terminal region of DFRP1 and DFRP2, as described earlier (Figures 2A and B). DFRP1 has a C-terminal region that wraps around DRG1, whilst this region in DFRP2 does not contact DRG2. We reasoned that this C-terminal region of DFRP1 may be preventing its association with ERAKY^DRG2^. Therefore, we created a V5-tagged DFRP1 vector in which the C-terminal residues 296-426 were removed, to test its effect on DRG binding. HA-DRG1/2 was transfected into HEK293T cells with or without co-transfection of either V5-tagged full length DFRP1 or the 1-295 truncation mutant, followed by an anti-HA IP and western blotting (Figure S7B). As mentioned earlier, this C-terminal region of DFRP1 is not required for the interaction with DRG1 (Ishikawa et al., 2005). Consistent with this previous finding, V5-DFRP1 1-295 retained the ability to interact with HA-DRG1 (Figure S7B). As expected, neither full length DFRP1 nor the DFRP1 1-295 truncation mutant interacted with wildtype HA-DRG2 (Figure S7B). Moreover, DFRP1 1-295 did not show any interaction with DRG2 ERAKY (Figure S7B), suggesting that this C-terminal region is not responsible for the lack of DFRP1 binding observed in Figure S7A.

### DFRP stimulated DRG GTPase activity is specific

Having successfully altered the specificity of DRG1, by switching its binding from DFRP1 to DFRP2 (Figure 6), we sought to investigate what functional consequences this change might elicit. DFRP1 is known to stimulate the GTPase activity of DRG1 by 3- to 4-fold, possibly by increasing the structural stability of DRG1 and its affinity for potassium ions (Francis et al., 2012; Pérez-Arellano et al., 2013). We therefore sought to investigate whether DFRP2 binding to DRG1 would have the same stimulatory affect. However, as it is not yet known whether DFRP2 is also capable of stimulating DRG2 GTPase activity, we investigated this first.

To investigate the potential for DFRP2 stimulated GTPase activity, we utilised a luminescence-coupled GTPase assay to measure the activity of Flag-tagged DRG2 protein purified from HEK293T cells, as reported previously (Westrip et al., 2023). We transfected and purified Flag-tagged wildtype DRG2 from HEK293T cells with and without co-transfection of DFRP2. In addition, we transfected and purified DRG2-Flag A260Y, as this mutation was earlier shown to disrupt DFRP2 binding (Figure 4D). Figure 8A shows the results of the GTPase assay and a representative Coomassie stained gel of purified DRG2-Flag protein. We also ran western blots to confirm the relative amount of bound DFRP2 (Figure 8B). The results indicate that loss of DFRP2 binding caused an equivalent decrease in the GTPase activity of DRG2 (Figures 8A and 8B). These results demonstrate that DFRP2 can stimulate the GTPase activity of DRG2, raising the possibility that DFRP2 might also promote DRG1 activity when in an engineered complex together.

**Figure 8:**
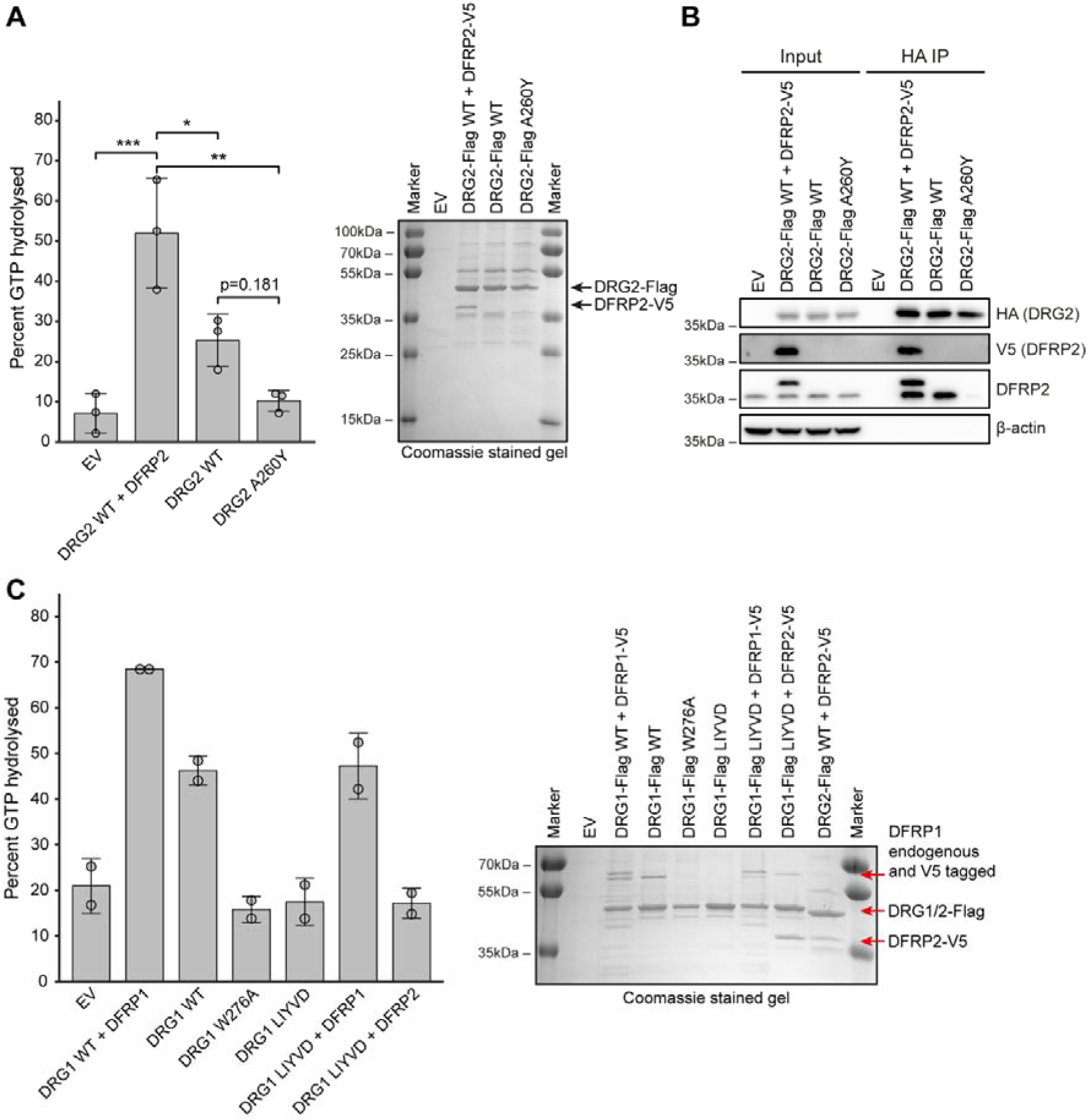
DFRPs stimulate specific DRG GTPase activity. **(A)** GTPase assay using DRG2-Flag purified from HEK293T cells using an anti-Flag immunoprecipitation (IP). The Coomassie stained gel (right) shows the purified protein used in the assay. Data represents the mean with error bars showing standard deviation. * = p<0.05. ** = p<0.01. *** = p<0.001. **(B)** Western blots using the purified protein from (A). **(C)** GTPase assay using purified DRG1-Flag protein shown in the Coomassie stained gel. Data represents the mean with error bars showing standard deviation. WT: wildtype. EV: empty vector plasmid.

We next tested the ability of DFRP2 to stimulate the activity of its engineered binding partner, DRG1 LIYVD, in an analogous approach to that described above. Flag-tagged wildtype or LIYVD^DRG1^ mutant DRG1 was transfected into HEK293T cells with either DFRP1-V5 or DFRP2-V5, followed by anti-Flag purification of the DRG/DFRP complexes. We also included the W276A^DRG1^ mutant as this was earlier shown to cause loss of DFRP1 binding (Figure S6D). As expected, loss of DFRP1 binding reduced the GTPase activity of DRG1 (Figure 8C) consistent with earlier reports in the literature (Francis et al., 2012; Pérez-Arellano et al., 2013). DRG1 LIYVD, on its own, showed reduced GTPase activity relative to wildtype DRG1 (Figure 8C). Interestingly, co-transfection of LIYVD^DRG1^ with DFRP1-V5 partially restored the loss of GTPase activity relative to wildtype DRG1-Flag co-transfected with DFRP1-V5. However, co-transfection with DFRP2-V5 did not increase the GTPase activity of DRG1 LIYVD, despite considerable DFRP2 binding (Figure 8C). Overall, these results suggest that whilst both DFRPs can stimulate GTPase activity, this function is specific for their respective DRG protein.

## Discussion

Here, we have built predicted structural models of the two human DRG/DFRP complexes using the AlphaFold2 algorithm (Jumper et al., 2021; Mirdita et al., 2022). With these structures, we have characterised the interactions between DRGs and DFRPs, including identifying specificity determining residues. In particular, we have identified five residues in DRG1 that are required for specifying DFRP1 binding over DFRP2. Mutating these five residues together in DRG1, switched binding from DFRP1 to DFRP2. Of these five positions, two residues (Y262^DRG1^ and D280^DRG1^) were especially important for specifying binding to DFRP1 over DFRP2. Mutating these DRG1 residues to their corresponding positions in DRG2 (Y262A^DRG1^ + D280Y^DRG1^) resulted in a partial rewiring of the specificity, in that DFRP2 binding was gained whilst DFRP1 binding was decreased (Figure 6). Both these residues likely function as positive and negative elements, in that they promote the DFRP1 interaction whilst also blocking DFRP2 binding.

The three other DRG1 mutations (L155E^DRG1^, I260R^DRG1^, and V264K^DRG1^) involved in the switch in binding formed part of a hydrophobic pocket that is occupied by the α6 helix of DFRP1. These mutations reduced the DFRP1 interaction, likely through steric clashes with the α6 helix, but are not expected to interact with DFRP2. Despite this, these DRG1 mutations contributed to the switch in binding from DFRP1 to DFRP2, presumably by inhibiting the DRG1-DFRP1 interaction and thus making DRG1 (within the context of the Y262A^DRG1^ + D280Y^DRG1^ mutations) more freely available for DFRP2 binding.

Protein binding specificity between paralogs has been well characterised in a number of examples, notably bacterial two-component signal transduction systems and toxin-antitoxin interactions (Aakre et al., 2015; Capra et al., 2010; Lite et al., 2020; Skerker et al., 2008). This has revealed that specificity is often controlled by only a handful of interface residues. Thus, it is perhaps unsurprising that so few mutations are needed in DRG1 in order to alter the DFRP binding specificity. However, it is interesting that DRG1 LIYVD still retained some ability to interact with DFRP1 (Figure 7). This suggests that the residues mutated in DRG1 play an important role in preventing DFRP2 binding and specifying the DFRP1 interaction but are not necessarily required for the DFRP1 interaction. This may be explained by the large size of the interface, which would indicate there are many other DRG1 residues responsible for the DFRP1 interaction. It may be possible to introduce further mutations to DRG1 LIYVD that could block the DFRP1 interaction.

In contrast to DRG1, mutating the same residues in DRG2 failed to produce any switch in binding, although the DFRP2 interaction was lost (Figure S7). There are several possible reasons for this. The interface is large and complicated and may therefore require many more mutations to switch the binding. Another possibility is that the specificity determining residues in DRG2 are not entirely located to the interface in question, and as such would have been missed by our focussed analysis. It is possible that residues in one part of a protein’s structure can affect residues in a distal location of a protein, through effects on the folding/packing of residues. As such, it is also appreciated that non-interface residues play a role in protein interactions, including the evolution of binding specificity (Ding et al., 2022). Future analysis of the DRG2/DFRP2 complex for non-interface residues that co-evolve with the interface may help in the identification of this type of specificity determining residue.

The large difference between DFRP1 and DFRP2 (albeit for the DFRP domain) suggests they contribute to the variation in function between DRG1 and DRG2. However, it still remains unclear as to whether DRG1 and DRG2 are functionally redundant without their specific DFRP interactions. If DFRP1 and DFRP2 are responsible for the difference in function between DRGs, then it may be possible for DRG1 and DRG2 to partially compensate for one another if their DFRP interactions were switched. In other words, the LIYVD^DRG1^:DFRP2 complex we have engineered in this paper may be able to functionally compensate for loss of DRG2, though this remains untested.

Both DRG/DFRP complexes have been implicated in diseases, including as potential oncogenes in cancer (Westrip et al., 2021). Protein-protein interactions have long been considered as potential targets for therapeutic intervention (Wells and McClendon, 2007). The work presented here may support the design of specific inhibitors, including, for example, those that target the hydrophobic pocket in the DRG1/DFRP1 complex. Further research into the mechanisms underlying DRG:DFRP interactions, and the functions of the resulting complexes, will aid our understanding of their physiological roles and their potential for clinical translation.

## Methods

### Cell culture

HEK293T cells were cultured in Dulbecco’s Modified Eagle Medium (DMEM, Gibco) containing 10% (v/v) foetal bovine serum (FBS, Sigma) and 100 international units/mL (IU/mL) penicillin and 100 µg/mL streptomycin (Gibco). Cells were grown in a humidified incubator set at 37°C, and 5% CO_2_.

### Co-IP experiments

HEK293T cells were seeded onto 6-well plates (4x10^5^ cells per well). The following day cells were transfected with plasmid DNA using FuGENE® 6 or FuGENE® HD as follows. For 1 well: 1µg of DNA and 3µl of FuGENE® (1:3 ratio) were combined with 100µl of Opti-MEM™ (Gibco), vortexed and incubated at room temperature for 30 minutes before adding to the cells. 48 hours after transfection the cells were harvested with 200µl of IP lysis buffer (100mM NaCl, 20mM TrisHCl [pH 7.4], 5mM MgCl_2_, 0.5% (v/v) NP-40) containing 1x SIGMAFAST™ protease inhibitor cocktail (Sigma-Aldrich, S8830). For each well/condition, 40µl of lysate was taken for input and the remaining lysate was combined with 20µl of either anti-HA agarose beads or anti-V5 agarose beads. The bead/lysate mix was left spinning overnight at 4°C. The beads were then washed 6 times in the IP lysis buffer, before beads were boiled at 95°C for 5 minutes in 20µl of 2x sample buffer. 6x sample buffer: 125mM Tris-HCl [pH6.8], 6% (w/v) SDS, 50% (v/v) glycerol, 225mM dithiotheitol (DTT), 0.1% (w/v) bromophenol blue. Sample were then analysed by western blotting, typically loading around 3-4µg of input sample and 25-30% of the IP sample.

### Endogenous IP

HEK293T cells were seeded onto 10cm plates (2.5x10^6^ cells/plate) and harvested in 2ml IP lysis buffer per 10cm plate. Primary antibody was then combined with 2-2.5mg of cell lysate and incubated overnight at 4°C. The following rabbit polyclonal antibodies were used: 5µg of anti-DFRP1 (Atlas Antibodies, HPA031099), 5µg of anti-DFRP2 (Genetex, GTX120331), 3µg of anti-DRG1 (ProteinTech, 13190-1-AP). For the control IP, an equivalent amount of anti-IgG antibody was used. For each IP condition, 60µL of Protein A beads (EMD Millipore, 16-156) were washed in IP lysis buffer then blocked for 1hour in 1%BSA in PBS, then washed again in IP lysis buffer. The beads were combined with the antibody/lysate mix and incubated for 4hours at 4°C, before being washed extensively in IP lysis buffer. The beads were then boiled in 60µl of 2x sample buffer at 95°C for 5 minutes.

### SDS-PAGE and western blotting

Proteins were resolved using sodium dodecyl sulphate-polyacrylamide gel electrophoresis (SDS-PAGE), specifically made in-house 12% polyacrylamide gels. Gels were run at 150 volts in1x Tris/Glycine/SDS (National diagnostics) running buffer. After electrophoresis, the protein was transferred to an Amersham™ Hybond™ P0.45 polyvinylidene difluoride (PVDF) membrane (GE Healthcare) in 1x Tris/Glycine transfer buffer (National diagnostics) containing 20% (v/v) methanol. The PVDF membrane was also activated in methanol prior to use. The transfer was run at a constant 320mA for 30 minutes for 1 gel or 50 minutes for 2 gels. After transfer, the membrane was blocked in 5% (w/v) milk powder dissolved in PBST (PBS with 0.1% (v/v) Tween-20) for 1 hour at room temperature. After blocking, the membrane was incubated in primary antibody overnight at 4°C or for 1-2 hours at room temperature for HRP linked antibodies. After primary antibody incubation, the membrane was washed in PBS-T, then incubated in the relevant horseradish peroxidase (HRP)-conjugated secondary antibody for 1 hour at room temperature. After further washing in PBS-T, the membrane was imaged using either the Clarity Western ECL reagent (Bio-Rad) or Femto (Thermo Scientific) in a Vilber Fusion Fx.

### GTPase assays

HEK293T cells were seeded onto 15cm plates (5x10^6^ cells/plate). The following day the cells were transfected with C-terminal Flag-tagged DRG1 or DRG2 (wildtype or mutant) with or without co-transfection of C-terminal V5-tagged DFRP1 of DFRP2. DNA was transfected using FuGENE® 6 or FuGENE® HD same as for co-IP experiments but scaled up to 10µg DNA per plate. 48hrs after transfection cells were harvested in IP lysis buffer. Flag-tagged protein was purified using 20µl anti-Flag® M2 magnetic beads (Sigma) per plate. After overnight incubation at 4°C the beads were washed in IP lysis buffer and protein was eluted in a GTPase buffer (100mM Tris pH8, 300mM KCl, 20mM MgCl_2_, 10% v/v glycerol) containing 0.15µg/ml Flag peptide.

GTPase activity of the purified protein was measure using the Promega GTPase-Glo™ kit (Promega, V7681), as per the manufacturer’s instructions. Assays were done on a 384-well plate, with three technical repeats for each DRG purification. For assaying DRG1, the reactions were incubated at 37°C for 18hrs. For assaying DRG2, the reactions were incubated at 37°C for 2hrs. Potassium (K^+^) was used at a final concentration of 150mM. The fluorescent output of the GTPase assays was measure using a PerkinElmer Enspire Multimode Plate reader.

### Structural and sequence analysis

Human DRG and DFRP sequences were downloaded from Uniprot and inputted into the online ColabFold server for structure prediction. All structural analysis was carried out using ChimeraX. Structure comparisons between predicted structures and already solved structure models were carried out using the Matchmaker tool. Interface residues (contacts between DRGs and DFRPs) were identified using the Find Contacts function with default parameters.

For sequence alignments, DRG and DFRP sequences (Tables 1 and 2) were identified using BLAST searches against the NCBI database. Sequences were aligned using the MUSCLE algorithm using MEGAX software. Shaded alignments with secondary structure were created using ESPript 3.0 software.

**Table 1:** DRG1/2 protein sequences.

| Protein | Organism | ID |
| --- | --- | --- |
| DRG1 | *Homo sapiens* | AAH20803.1 |
| DRG1 | *Xenopus leavis* | NP_001084013.1 |
| DRG1 | *Drosophila melanogaster* | NP_536733.1 |
| DRG1 | *Saccharomyces cerevisiae* | AJP36937.1 |
| DRG1 | *Arabidopsis thaliana* | NP_195662.1 |
| DRG2 | *Homo sapiens* | NP_001379.1 |
| DRG2 | *Xenopus leavis* | NP_001079983.1 |
| DRG2 | *Drosophila melanogaster* | NP_650822.1 |
| DRG2 | *Saccharomyces cerevisiae* | AJR85197.1 |
| DRG2 | *Arabidopsis thaliana* | NP_001077555.1 |

**Table 2:** DFRP1/2 protein sequences.

| Protein | Organism | ID |
| --- | --- | --- |
| DFRP1 | *Homo sapiens* | Q8WU90 |
| DFRP1 | *Xenopus leavis* | NP_001086961.1 |
| DFRP1 | *Drosophila melanogaster* | NP_001286202.1 |
| DFRP1 | *Saccharomyces cerevisiae* | AJU02303.1 |
| DFRP1 | *Arabidopsis thaliana* | NP_179618.1 |
| DFRP2 | *Homo sapiens* | Q9H446 |
| DFRP2 | *Xenopus leavis* | NP_001087542.1 |
| DFRP2 | *Drosophila melanogaster* | NP_001262901.1 |
| DFRP2 | *Saccharomyces cerevisiae* | KZV12390.1 |
| DFRP2 | *Arabidopsis thaliana* | NP_175584.1 |

## Acknowledgments

M.L.C is supported by a CRUK Programme Foundation Award (C33483/A2567).

## Author Contributions

C.A.E.W conceived and designed the project, carried out the experiments, analysed the data, and drafted the manuscript. M.L.C analysed data, supervised and funded the project, and revised the manuscript. S.J.S advised on structural analyses and manuscript preparation.

