## Supplemental Information for "Rewiring protein binding specificity in paralogous DRG/DFRP complexes"

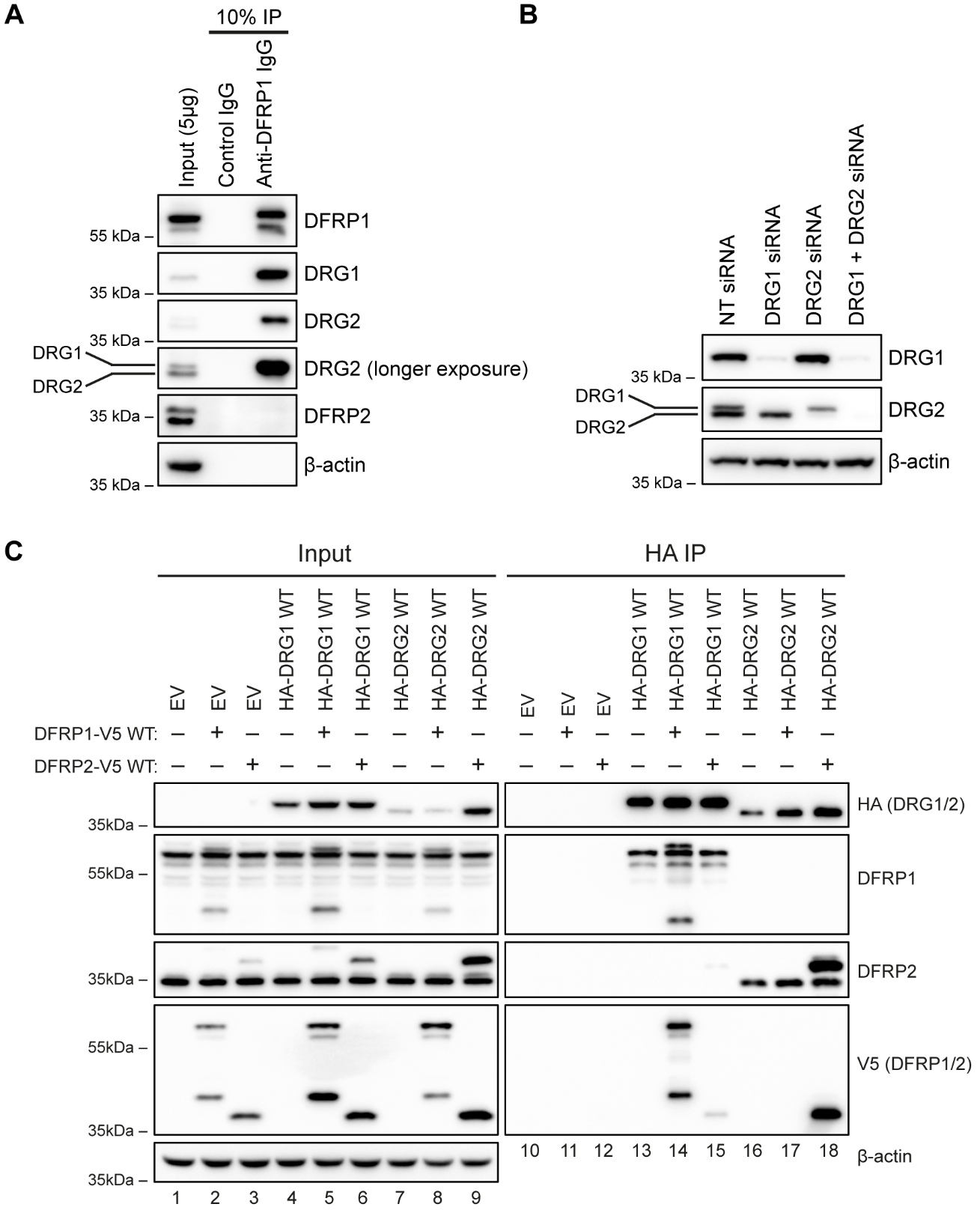

**Figure S1: DRG-DFRP complexes. (A)** Endogenous immunoprecipitation (IP) of DFRP1 from HEK293T cells. Note that anti-DRG2 antibodies detect both DRG1 and DRG2 (B), and therefore the slightly slower migrating band in the IP sample for the DRG2 blot is likely DRG1. **(B)** Cross-reactivity between anti-DRG2 antibodies and the DRG1 protein. HEK293T cells were transfected with siRNA targeting DRG1 or DRG2 for 72 hours followed by western blotting. NT: non-targeting siRNA. Note that the anti-DRG2 antibody detects two bands, the upper one of the two being reduced by DRG1 siRNA. **(C)** Specific binding between DRGs and DFRPs. HEK293T cells were transfected for 48 hours with HA-DRG1 or HA-DRG2 with and without DFRP1-V5 or DFRP2-V5. Cells were harvested and lysates used in an anti-HA IP before western blotting. Lane 15 demonstrates the weak binding observed between DRG1 and DFRP2 when both proteins are overexpressed. EV: empty vector plasmid. WT: wildtype.

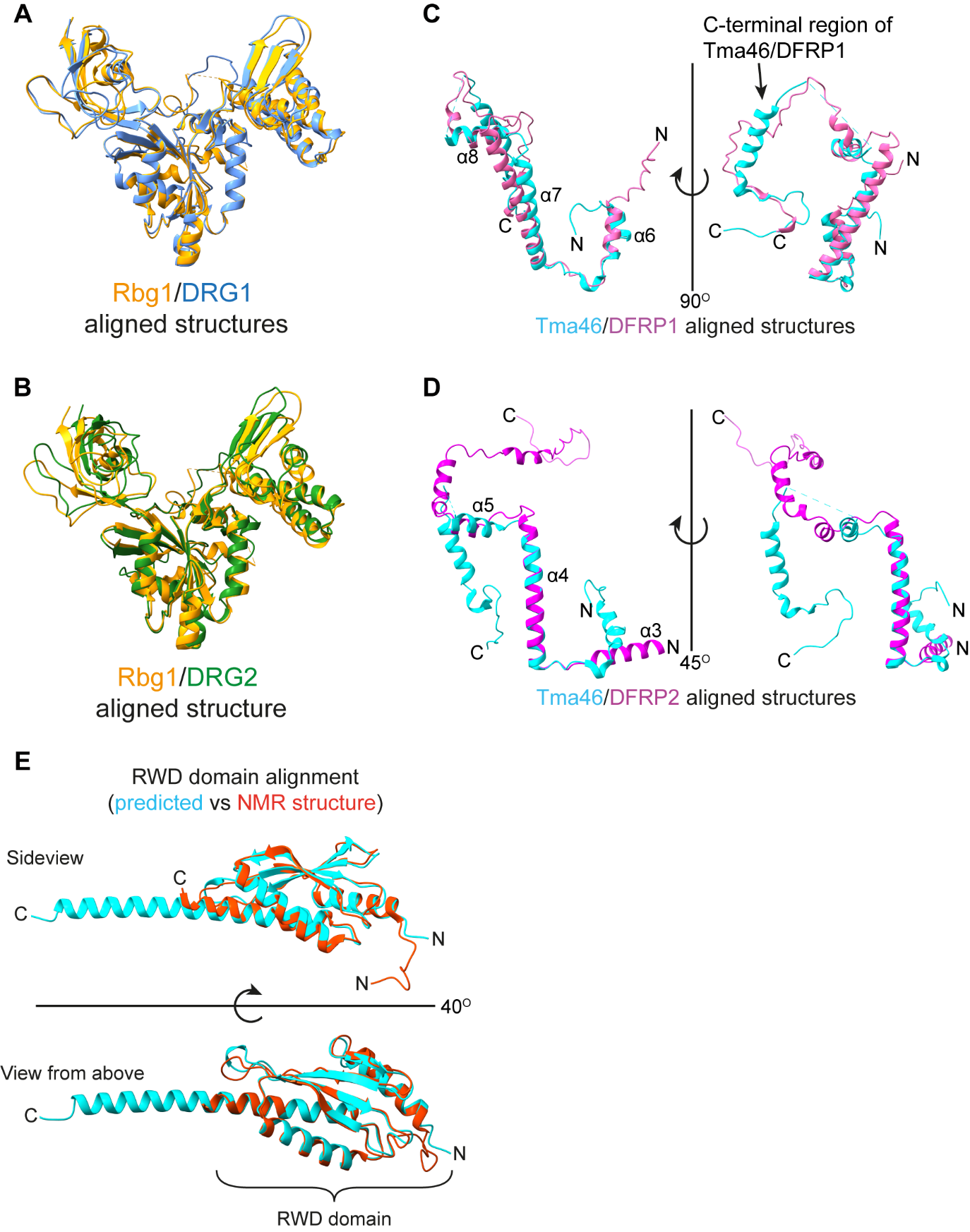

**Figure S2: Comparison of AlphaFold predicted models and crystal/NMR structure. (A)** Structural alignment of Rbg1 (Pdb: 4A9A) (orange) with AlphaFold predicted DRG1 structure (blue). RMSD between 229 pruned atom pairs is 1.04 Å; (across all 355 pairs: 2.96 Å). **(B)** Structural alignment of Rbg1 (Pdb:4A9A) (orange) with predicted DRG2 structure (green). RMSD between 197 pruned atom pairs is 1.18 Å; (across all 353 pairs: 3.19 Å). **(C)** Structural alignment of predicted DFRP1 (K226-R340) and Tma46 (Pdb: 49A9). RMSD (Root mean square deviation) between 37 pruned atom pairs is 1.09 angstroms (Å); (across all 95 pairs: 17.71 Å). **(D)** Structural alignment of Tma46 (Pdb: 4A9A) with the predicted DFRP2 structure (Q128-D243). RMSD between 28 pruned atom pairs is 0.54 Å; (across all 88 pairs: 37.84 Å). For (A-E) structural alignments and RMSD calculations were done in ChimeraX using the Matchmaker tool. **(E)** Structural alignment of the RWD domain from the predicted DFRP2 structure (coloured cyan) with the already reported NMR structure of the RWD domain (Pdb: 2EBM) (coloured red). RMSD between 114 pruned atom pairs is 0.90 Å; (across all 121 pairs: 1.13 Å).

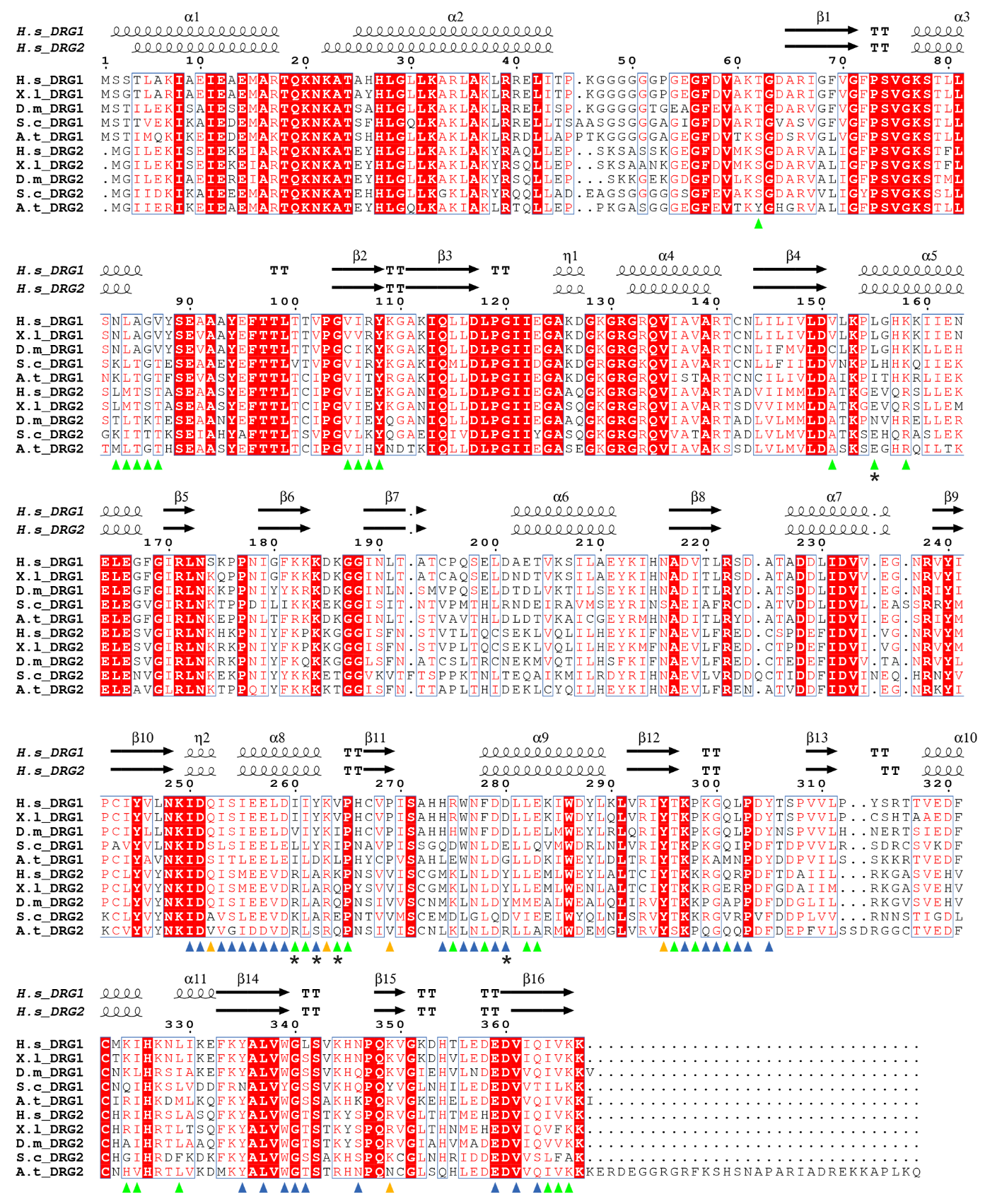

**Figure S3: DRG1 and DRG2 protein sequence alignment showing interface residues.** Alignment of DRG1 and DRG2 protein sequences created using the MUSCLE algorithm in MEGA11. The final shaded alignment was produced using ESPript 3.0. The secondary structure from the predicted DRGs in Figures 2A and 2B is shown. The triangles indicate the positions in DRG1/2 that contact DFRP1/2. Specifically, the blue triangles show residues that are in both interfaces, whilst green represents DRG1/DFRP1 only. The orange triangles show DRG2/DFRP2 specific interface residues. The asterisks indicate positions we identified as potentially having a role in binding specificity. The numbers above the alignment correspond to the DRG1 amino acid position.

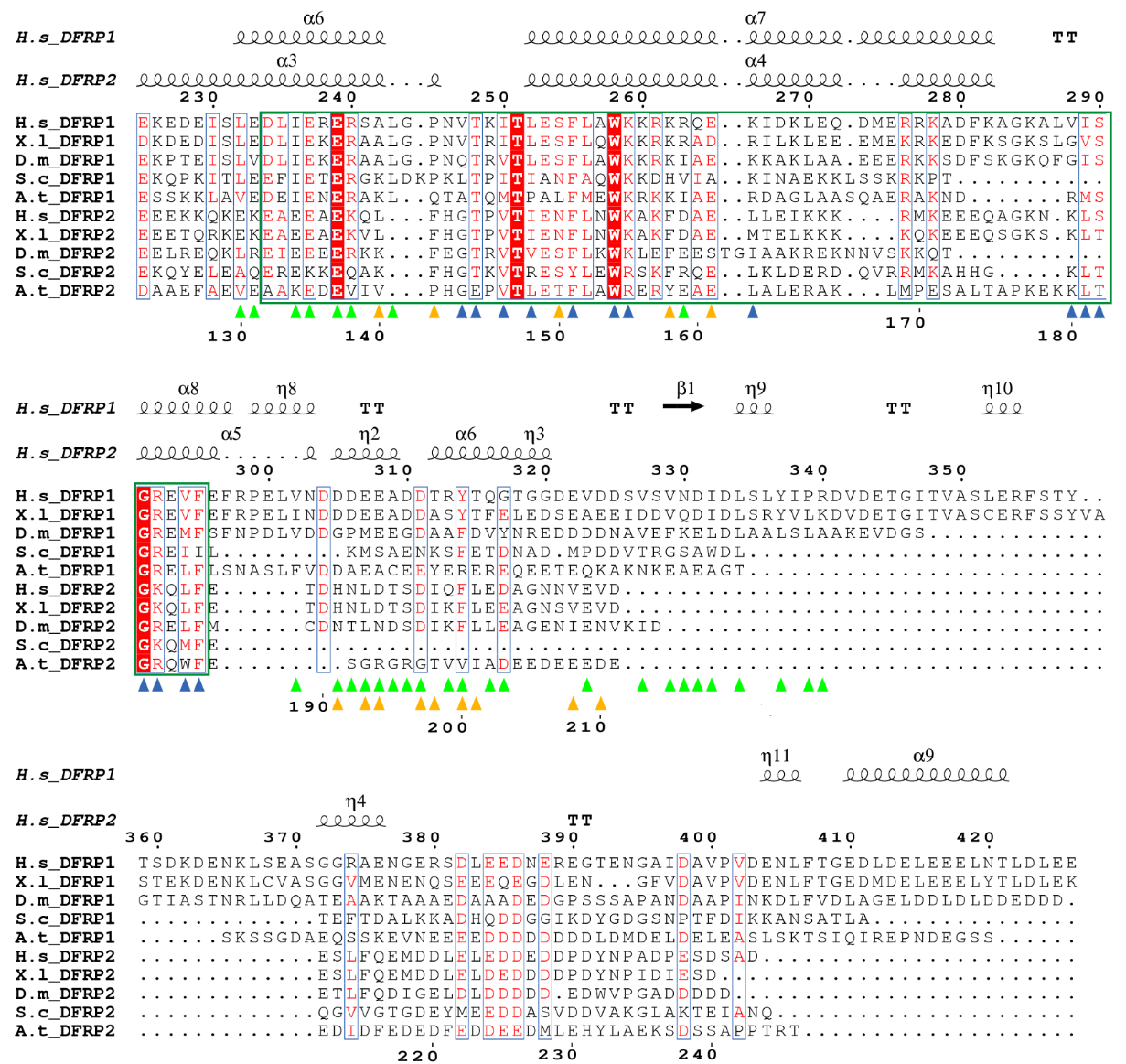

**Figure S4: DFRP1/2 protein sequence alignment.** DFRP1 and DFRP2 proteins sequences were aligned using the MUSCLE algorithm in MEGA11. The final shaded alignment was produced using ESPript 3.0. Secondary structure from the predicted DFRPs is shown. The blue triangles show residues that are in both interfaces, whilst green represents DRG1/DFRP1 only and orange shows DRG2/DFRP2 specific interface residues. The DFRP domain is indicated by the green box. The region 300-340 of DFRP1 is not conserved with DFRP2, so interface residues have been labelled separately. The number above the sequences corresponds to the human DFRP1 amino acid position, whilst numbers below indicate the human DFRP2 positions.

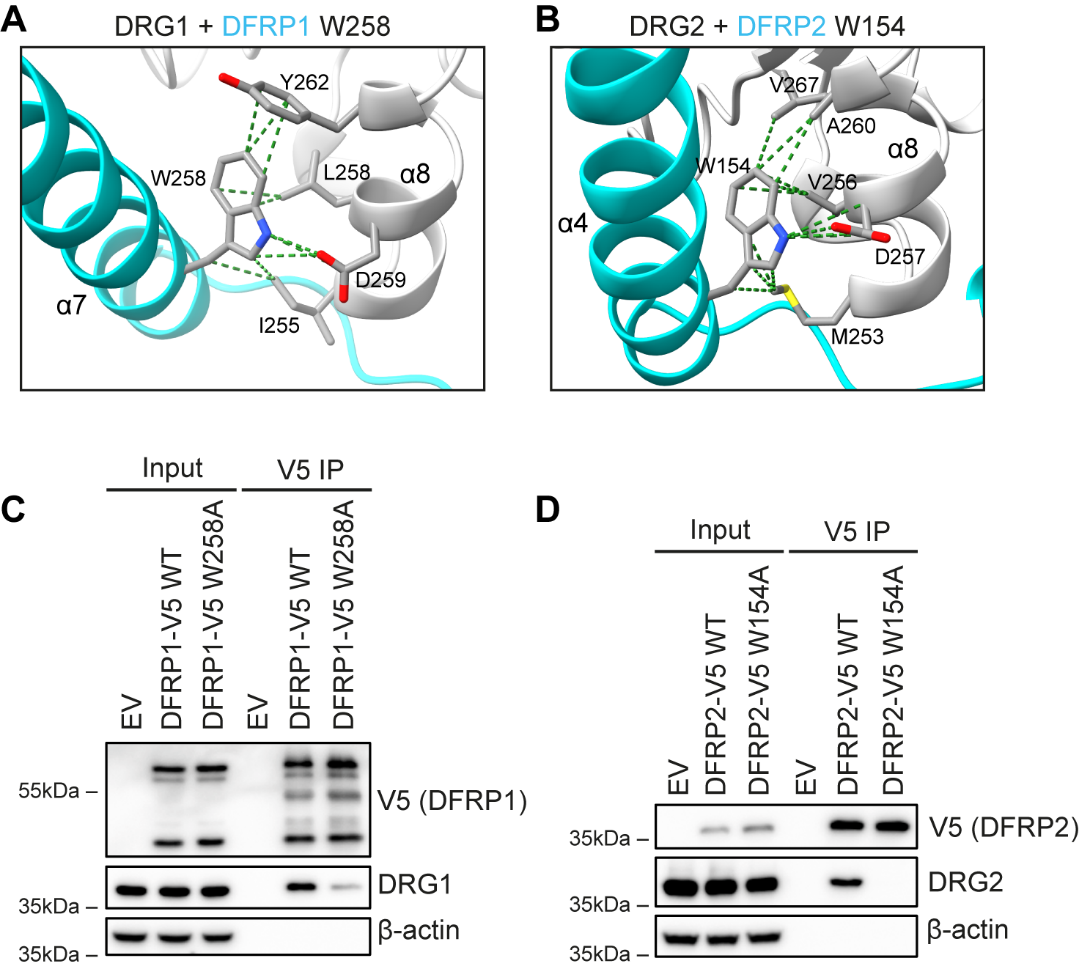

**Figure S5:** **Conserved DFRP1/2 tryptophan is important for DRG1/2 binding. (A)** Structural analysis of W258^DFRP1^ in the predicted DRG1/DFRP1 structure, revealing several contacts made with DRG1. DRG1 is grey, and DFRP1 is coloured cyan. **(B)** Structural analysis of W154DFRP2 in the predicted DRG2/DFRP2 structure, showing contacts made with DRG2. DRG2 and DFRP2 are coloured grey and cyan respectively. **(C)** HEK293T cells were transfected with a vector encoding C-terminally V5-tagged DFRP1 wildtype (WT) or the W258A mutation, followed by an anti-V5 immunoprecipitation (IP) and western blotting. **(D)** Same as in (C) but with C-terminally V5-tagged DFRP2 WT or the W154A mutation. EV: empty vector plasmid.

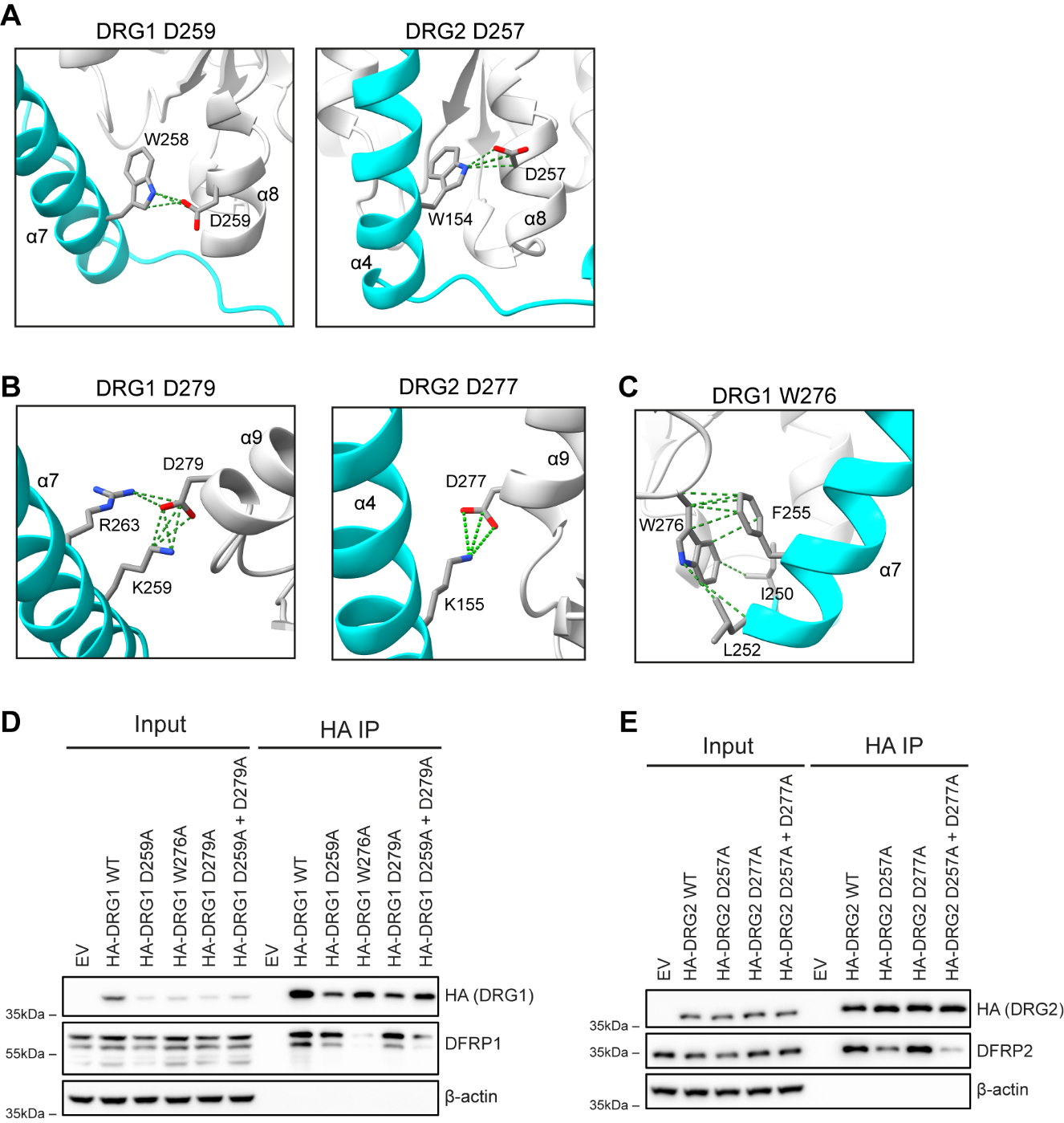

**Figure S6: DRG1 and DRG2 residues important for DFRP binding. (A)** Structural analysis of D259^DRG1^ (left panel) and D257^DRG2^ (right panel), showing both residues contact the invariant DFRP domain tryptophan (W258^DFRP1^ and W154^DFRP2^, respectively). **(B)** Structural analysis of D279^DRG1^ (left panel) and D277^DRG2^ (right panel) showing both residues contact positively charged residues in DFRP1 and DFRP2, respectively. **(C)** Structural analysis of W276^DRG1^ and the pi-pi interactions it may form with F255^DFRP1^. **(D)** HEK293T cells were transfected with vectors encoding HA-DRG1 wildtype (WT) or the indicated mutations, followed by an anti-HA immunoprecipitation (IP) and western blotting. **(E)** Same as in (D) except vectors encoded HA-DRG2 WT or mutants. EV: empty vector plasmid.

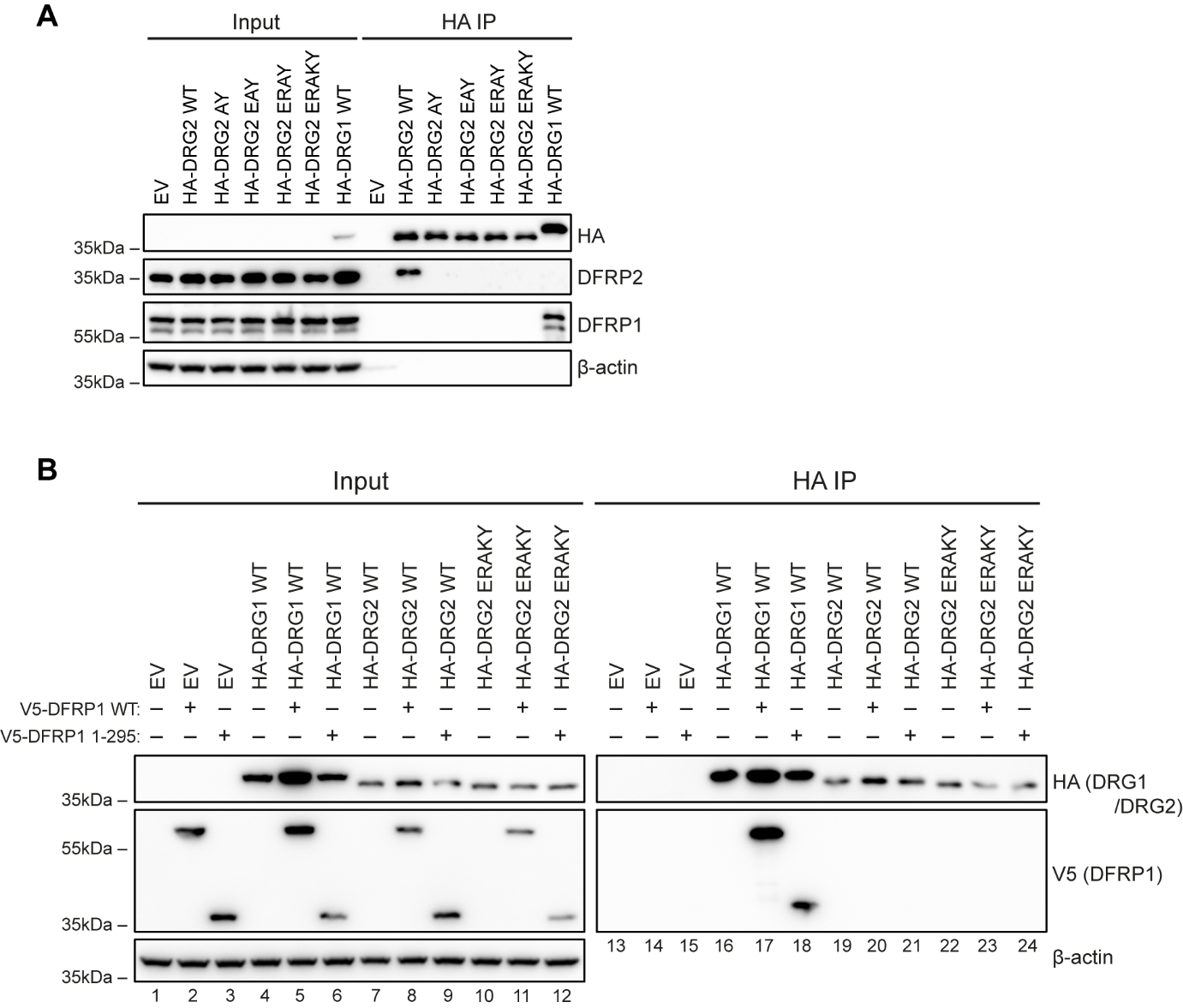

**Figure S7: Combined DRG2 mutations do not switch DFRP binding specificity. (A)** HEK293T cells were transfected with vectors encoding HA-DRG2 wildtype (WT) or combinations of the following five mutations indicated by their amino acid position in DRG2 (E153L^DRG2^, R258I^DRG2^, A260Y^DRG2^, K262V^DRG2^ and Y278D^DRG2^). The lysate was used in an anti-HA immunoprecipitation (IP) followed by western blotting. **(B)** HEK293T cells were transfected with vectors encoding HA-DRG1, HA-DRG2 WT or HA-DRG2 ERAKY, with or without co-transfection of full length V5-DFRP1 WT or a truncation mutant V5-DFRP1 1-295. Lysates were subjected to an anti-HA IP followed by western blotting. EV: empty vector plasmid.
